# Generation and characterisation of *P. falciparum* parasites with a G358S mutation in the PfATP4 Na^+^ pump and clinically relevant levels of resistance to some PfATP4 inhibitors

**DOI:** 10.1101/2022.01.11.475938

**Authors:** Deyun Qiu, Jinxin V. Pei, James E. O. Rosling, Dongdi Li, Yi Xue, Jocelyn Sietsma Penington, Krittikorn Kümpornsin, Yi Tong Vincent Aw, Jessica Yi Han Aw, Heath Hasemer, Adelaide S. M. Dennis, Melanie C. Ridgway, Anthony T. Papenfuss, Marcus C. S. Lee, Giel G. van Dooren, Kiaran Kirk, Adele M. Lehane

**Affiliations:** Research School of Biology, Australian National University, Canberra, ACT 2600, Australia; Bioinformatic Division, The Walter & Eliza Hall Institute of Medical Research, Parkville, VIC 3052, Australia; Wellcome Sanger Institute, Wellcome Genome Campus, Hinxton CB10 1SA, United Kingdom; Department of Medical Biology, The University of Melbourne, Parkville, VIC 3052, Australia

## Abstract

Small-molecule inhibitors of PfATP4, a *Plasmodium falciparum* protein that is believed to pump Na^+^ out of the parasite while importing H^+^, are on track to become much-needed new antimalarial drugs. The spiroindolone cipargamin is poised to become the first PfATP4 inhibitor to reach the field, having performed strongly in Phase 1 and 2 clinical trials. Previous attempts to generate cipargamin-resistant parasites in the laboratory have yielded parasites with reduced susceptibility to the drug; however, the highest 50% inhibitory concentration reported to date is 24 nM. Here, we show that *P. falciparum* parasites can acquire a clinically-significant level of resistance to cipargamin that enables them to withstand micromolar concentrations of the drug. Independent experiments to generate high-level cipargamin resistance using different protocols and strains led to the same change each time – a G358S mutation in PfATP4. Parasites with this mutation showed high-level resistance not only to cipargamin, but also to the dihydroisoquinolone (+)-SJ733. However, for certain other (less clinically advanced) PfATP4-associated compounds the G358S mutation in PfATP4 conferred only moderate resistance or no resistance. The G358S mutation in PfATP4 did not affect parasite susceptibility to antimalarials that do not target PfATP4. The G358S mutation in PfATP4, and the equivalent mutation in the *Toxoplasma gondii* ATP4 homologue (G419S), decreased the sensitivity of the Na^+^-ATPase activity of ATP4 to inhibition by cipargamin and (+)-SJ733, and decreased the sensitivity of parasites expressing these ATP4 mutations to disruption of parasite Na^+^ regulation by cipargamin- and (+)-SJ733. The G358S mutation in PfATP4 reduced the affinity of the protein for Na^+^ and was associated with an increase in the parasite’s resting cytosolic Na^+^ concentration; however, no significant defect in parasite growth rate was observed. Our findings suggest that codon 358 in *pfatp4* should be monitored closely in the field as a molecular marker for cipargamin resistance, and that PfATP4 inhibitors in clinical development should be tested for their activity against PfATP4^G358S^ parasites.

## Introduction

Malaria, an ancient mosquito-borne disease caused by *Plasmodium* parasites, killed an estimated 627,000 people in 2020, of whom approximately 500,000 were children under the age of five years (1). Among the *Plasmodium* species that cause disease in humans, *Plasmodium falciparum* causes the vast majority of deaths (1). *P. falciparum* parasites resistant to most available antimalarials are widespread, and parasites with reduced susceptibility to the latest first-line malaria treatments, which consist of an artemisinin derivative paired with a quinoline-related compound, are spreading (2,3). The situation is precarious, as the widespread failure of artemisinin-based combination therapies would render malaria more difficult to treat (4,5). Thus, to safeguard malaria treatment, the development of novel antimalarial structures with new modes of action is required, along with strategies to protect them from resistance.

In the last 12 years millions of compounds have been screened for their ability to kill asexual blood-stage *P. falciparum* parasites (6-8). This has led to the discovery of promising new antimalarial drug classes, including the spiroindolones (9). In an attempt to identify the target of the spiroindolones, *in vitro* evolution experiments were performed in which *P. falciparum* parasites were exposed to spiroindolones until parasites displaying low-level resistance emerged. This resistance was associated with mutations in the gene encoding the P-type ATPase PfATP4, which was localised to the parasite plasma membrane (9). PfATP4 was originally thought to be a Ca^2+^ transporter (10); however Spillman *et al*. (11) provided evidence that it is in fact a Na^+^ transporter that exports Na^+^ from the parasite cytosol, whilst importing H^+^ equivalents. Spiroindolones cause a rapid increase in the [Na^+^] in the parasite cytosol ([Na^+^]_cyt_) (11). Parasites with mutations in PfATP4, showing low-level resistance to spiroindolones, were found to be less sensitive to the Na^+^-dysregulating effects of spiroindolones than their parents, and to have a higher resting [Na^+^]_cyt_ (11). These observations are consistent with the resistance-conferring mutations impacting on the (Na^+^-efflux) function of PfATP4 (11).

As well as causing a rise in [Na^+^]_cyt_ (and thereby dissipating the inward [Na^+^] gradient across the parasite plasma membrane), spiroindolones cause a variety of other physiological perturbations including: an alkalinisation of the parasite cytosol, which *increases* the pH gradient across the parasite plasma membrane (11); an increase in the volume of parasites and parasitised erythrocytes, attributable to the osmotic consequences of the [Na^+^]_cyt_ increase (12); a reduction in cholesterol extrusion from the parasite plasma membrane resulting from the increase in [Na^+^]_cyt_ (13); and an increase in the rigidity of erythrocytes infected with ring-stage parasites (14). Spiroindolones have also been shown to inhibit a Na^+^-dependent, pH-sensitive ATPase in parasite membrane preparations (11,15). Together, the available data are consistent with PfATP4 functioning as an ATP-dependent transporter on the parasite plasma membrane, extruding Na^+^ from the parasite while importing H^+^.

In contrast to *P. falciparum* parasites, which experience a high external Na^+^ concentration for the majority of their 48 h asexual life cycle (16) and for which a reduction of *pfatp4* expression is deleterious to growth (17), *T. gondii* parasites are only known to be exposed to a high external Na^+^ concentration for the brief period in their lytic cycle in which they are extracellular, and can survive and proliferate when their ATP4 homologue (TgATP4) is not expressed (18). Physiological studies with *T. gondii* parasites lacking TgATP4 expression have provided further evidence that ATP4 proteins export Na^+^ while importing H^+^ (18).

In the years since the discovery of the spiroindolones, a large number of chemically diverse compounds have been found to perturb parasite physiology in the same manner as the spiroindolones (19-25). These include the dihydroisoquinolone (+)-SJ733 (23), which has recently been tested in humans (26), the pyrazoleamide PA21A050 (25), and numerous less clinically-advanced compounds (19-22,24). The spiroindolone cipargamin (previously referred to as KAE609 and NITD609) is the most clinically-advanced of the compounds proposed to target PfATP4.

Oral formulations of cipargamin have been tested in multiple Phase 1 and Phase 2 clinical trials, with testing of an intravenous formulation suitable for the treatment of severe malaria underway (reviewed in (27)). Among the important attributes of cipargamin are (i) its ability to achieve rapid clearance of both *P. falciparum* and *P. vivax* parasites with minimal side effects (28), (ii) its favourable pharmacokinetic properties, which may make it a suitable component of a single-dose therapy (27), and (iii) its activity against sexual stage parasites in laboratory studies, which may translate into transmission-blocking activity in the field (29-31).

Another important consideration for compounds in clinical development is the risk of resistance developing (32). There is significant variability in the frequency with which *P. falciparum* parasites acquire resistance to different compounds. For example, atovaquone resistance occurs at a high frequency of ∼ 10^−5^ under certain conditions in some strains (i.e. 1 in 10^5^ parasites acquire resistance) (33), whereas for other compounds it has not yet been possible to generate resistance in the laboratory. It has been known since 2010 that long-term drug exposure experiments with cipargamin yield PfATP4-mutant parasites with low-level resistance (9). The frequency with which parasites acquired low-level resistance when exposed continuously to 2.5 nM cipargamin varied from ∼ 2 × 10^−8^ to 1 × 10^−7^, depending on the genetic background of the strain (34). *In vitro* evolution experiments have also been performed with other spiroindolones and with numerous chemically distinct compounds that also display the physiological hallmarks of PfATP4 inhibition, yielding resistant parasites with mutations in *pfatp4* in each case (20,21,23-25). More than 40 different resistance-associated SNPs have been reported in *pfatp4*, with most individual parasite lines having a single mutation in PfATP4, a small number having two mutations, and one reported to have three mutations (9,20,21,23-25,34).

It has been suggested that the frequency of resistance to PfATP4 inhibitors may be lower in the field than it is *in vitro*, as a result of the rapid speed by which the inhibitors kill parasites *in vivo* and fitness costs associated with resistance (23). *In vitro* growth competition experiments with asexual parasites have been reported for two (+)-SJ733-resistant PfATP4-mutant *P. falciparum* lines (PfATP4^L350H^ and PfATP4^P996T^), and both of these mutants were found to have a growth disadvantage relative to their parents (23). However, in a different study, PfATP4 mutant parasites selected for resistance to the aminopyrazole GNF-Pf4492 (PfATP4^A211T^, PfATP4^I203L/P990R^ and PfATP4^A187V^) were reported to grow at the same rate as their parents (20). Thus, it appears that some resistance-conferring PfATP4 mutations confer a growth disadvantage while others do not.

To date, the level of resistance that has been achieved *in vitro* for cipargamin has been modest. At least 18 parasite lines with different *pfatp4* mutations have been reported after *in vitro* evolution experiments with cipargamin (9,34). The 50% inhibitory concentrations (IC_50_s) for cipargamin against parasite growth ranged from 1.5 – 24.3 nM for the mutant parasites, compared to 0.4 – 1.1 nM for the parental lines used in these experiments. If parasites with a similar level of resistance to the *in vitro* selected parasites were to emerge in the field, it is likely that cipargamin would remain effective against them. In clinical trials, cipargamin has been found to reach supramicromolar concentrations in human plasma for sustained periods (28,35-37).

In this study, we aimed to determine whether parasites are able to acquire a clinically significant level of resistance to cipargamin, and if so, to investigate the mechanism involved and its effect on parasite physiology, growth rate, and susceptibility to other PfATP4 inhibitors and unrelated antimalarials.

## Results

### Generation of *P. falciparum* parasites highly resistant to cipargamin and other PfATP4 inhibitors

#### Continued exposure of parasites with low-level cipargamin resistance to the drug

To determine whether high-level resistance to cipargamin is achievable by *P. falciparum* parasites, we exposed two independent parasite cultures to incrementally increasing concentrations of cipargamin over the course of four months (**Fig. 1**). We commenced this experiment with a parasite line that had been selected for cipargamin resistance previously (NITD609-R^Dd2^ clone#2 (9); referred to here as Dd2-PfATP4^T418N,P990R^). This line displays low-level resistance to cipargamin (**Table 1**) and has two mutations (T418N and P990R) that are not present in its Dd2 strain parent.

**Table 1.** Susceptibility of the *P. falciparum* lines generated through *in vitro* evolution in this study and their parents to antiplasmodial compounds. The IC_50_ values (mean ± SEM) for growth inhibition for the parasite lines and compounds indicated are shown, with the number of independent experiments shown in brackets. Within each parasite set, all lines were tested in parallel in each experiment. For each compound, the IC_50_ values for each cipargamin-resistant line were compared with those of its direct parent (Dd2 for Dd2-PfATP4^T418N,P990R^; Dd2-PfATP4^T418N,P990R^ for HCR1 and HCR2; and Dd2-Polδ for Dd2-Polδ-PfATP4^G358S^) using paired t-tests.

|  | Mean IC<sub>50</sub> value ± SEM in nM |  |  |  |  |  |
| --- | --- | --- | --- | --- | --- | --- |
|  | Set 1 |  |  |  | Set 2 |  |
|  | Dd2 | Dd2-PfATP4<sup>T418N,P990R</sup> | HCR1 | HCR2 | Dd2-Polδ | Dd2-Polδ-PfATP4<sup>G358S</sup> |
| Cipargamin | 0.68 ± 0.03 (16) | 9.17 ± 0.46 (16)<br>***P < 0.001 | 2840 ± 190 (16)<br>***P < 0.001 | 2920 ± 200 (16)<br>***P < 0.001 | 1.43 ± 0.20 (7) | 1420 ± 100 (7)<br>***P < 0.001 |
| (+)-SJ733 | 60 ± 2 (5) | 331 ± 11 (5)<br>***P < 0.001 | 21500 ± 2400 (5)<br>***P < 0.001 | 17600 ± 2000 (5)<br>***P = 0.001 | 40.7 ± 4.4 (4) | 9070 ± 1010 (4)<br>**P = 0.003 |
| PA21A050 | 1.20 ± 0.04 (3) | 2.70 ± 0.16 (3)<br>*P = 0.01 | 61 ± 10 (3)<br>*P = 0.03 | 60 ± 4 (3)<br>**P = 0.005 | 2.42 ± 0.24 (4) | 12.9 ± 1.2 (4)<br>**P = 0.002 |
| Chloroquine | 127 ± 21 (4) | 131 ± 11 (4)<br>P = 0.7 | 115 ± 16 (4)<br>P = 0.06 | 167 ± 21 (4)<br>*P = 0.03 | 185 ± 35 (5) | 182 ± 27 (5)<br>P = 0.8 |
| Dihydro-artemisinin | 2.17 ± 0.28 (4) | 2.44 ± 0.23 (4)<br>P = 0.1 | 1.55 ± 0.21 (4)<br>**P = 0.005 | 1.51 ± 0.12 (4)<br>*P = 0.05 | 2.15 ± 0.21 (4) | 2.58 ± 0.27 (4)<br>P = 0.4 |
| MMV006656 | Not tested |  |  |  | 344 ± 43 (4) | 1800 ± 230 (4)<br>*P = 0.01 |
| MMV665949 | Not tested |  |  |  | 4270 ± 620 (4) | 4870 ± 880 (4)<br>P = 0.4 |

**Fig. 1.**
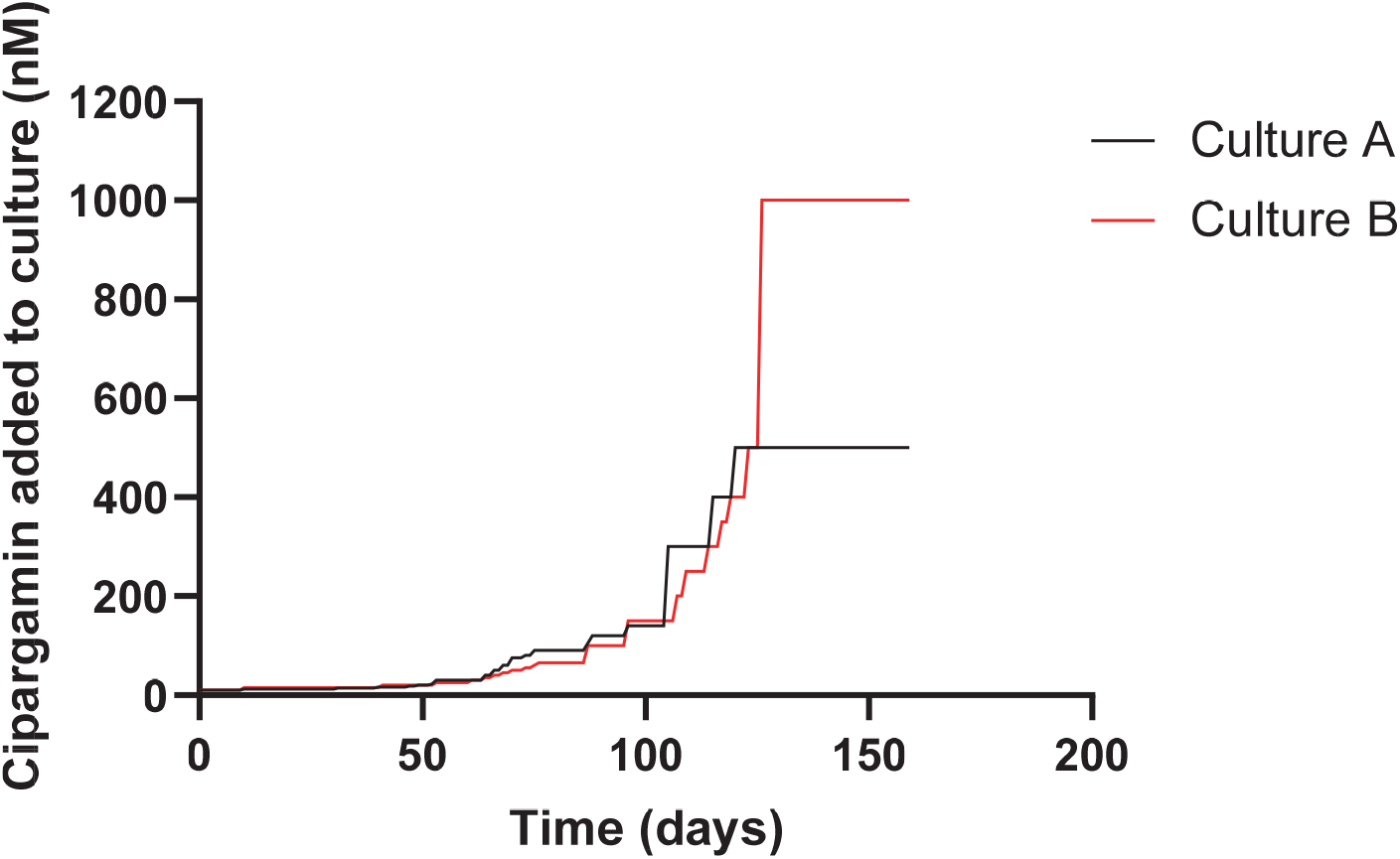
Generation of parasites with high-level cipargamin resistance from Dd2-PfATP4^T418N,P990R^ parasites having low-level resistance. Dd2-PfATP4^T418N,P990R^ parasites were exposed to incrementally increasing concentrations of cipargamin. Two independent selections were performed (shown in red and black).

At the end of the drug exposure period, parasites from both cultures were highly resistant to cipargamin, with IC_50_ values of 5.9 ± 0.2 μM (culture A; mean ± SEM, n = 3) and 6.2 ± 0.2 μM (culture B; mean ± SEM, n = 3). By comparison, the initial Dd2-PfATP4^T418N,P990R^ parasite line had an IC_50_ value for cipargamin below 10 nM and its Dd2 parent had a subnanomolar IC_50_ value (**Table 1**). We subjected the cipargamin-selected cultures to limiting dilution and selected clonal lines (‘highly-cipargamin-resistant (HCR)’ clones 1 and 2, from culture B) for further characterisation. The HCR1 and HCR2 parasites were highly resistant to cipargamin, having IC_50_ values 315 - 326 fold higher than that of their Dd2-PfATP4^T418N,P990R^ parents, and 4200 - 4280 fold higher than that of wild-type Dd2 parasites (mean, n = 16; **Fig. 2**; **Table 1**).

**Fig. 2.**
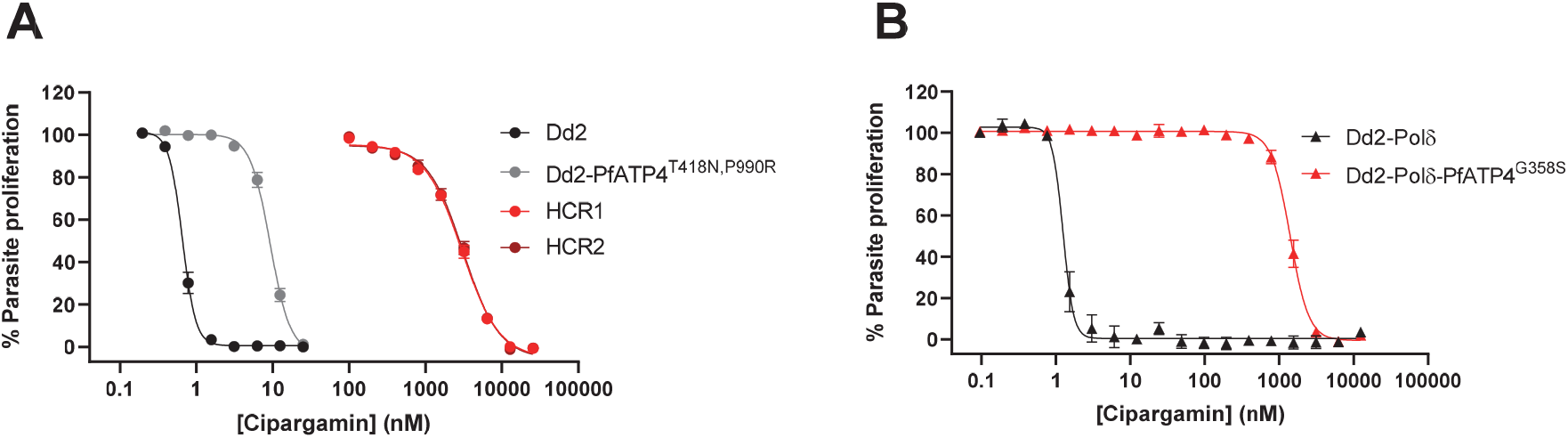
HCR and Dd2-Polδ-PfATP4^G358S^ parasites display a high level of resistance to cipargamin. **A**. Inhibition of parasite proliferation by cipargamin in Dd2 parasites (black circles), Dd2-PfATP4^T418N,P990R^ parasites (grey circles), and highly-cipargamin-resistant (HCR) parasites (clone #1 in red circles and clone #2 in dark red circles). **B**. Inhibition of parasite proliferation by cipargamin in Dd2-Polδ parasites (black triangles) and Dd2-Polδ-PfATP4^G358S^ parasites (red triangles). The data shown are the mean (± SEM) from 16 independent experiments for **A**, and from seven independent experiments for **B**, each performed on different days. All lines for which data are shown on the same panel were tested in parallel. Where not shown, error bars fall within the symbols.

We also attempted to generate highly cipargamin-resistant parasites using a single-step protocol in which parasites were exposed to a single high concentration of cipargamin. We exposed two 150 mL cultures, each containing ∼ 6.6 × 10^9^ Dd2-PfATP4^T418N,P990R^ parasites, to a ∼ 10 × IC_50_ concentration of cipargamin (102 nM). We did not recover viable resistant parasites using this approach with Dd2-PfATP4^T418N,P990R^ parasites.

#### Single step cipargamin selection with ‘hypermutator’ parasites

We then attempted to generate highly cipargamin resistant parasites using a single step procedure with a *P. falciparum* Dd2 line with a hypermutator phenotype. These parasites, referred to here as ‘Dd2-Polδ’ parasites, were modified using CRISPR-Cas9 to introduce two mutations into the gene encoding the DNA polymerase delta subunit in order to impair the protein’s proofreading function (Kümpornsin *et al*., publication in preparation). Dd2-Polδ parasites (three independent cultures, each containing 1.5 × 10^8^ parasites) were exposed to cipargamin at a single concentration of 25 nM, with one further culture containing 1.5 × 10^8^ parasites exposed to 100 nM cipargamin. Viable parasites were observed in one of the cultures containing 25 nM cipargamin (‘cipargamin-selected Culture 1’) after 17 days. Resistant parasites did not emerge in the other three cultures within 25 days and these cultures were discarded. A clonal line from cipargamin-selected Culture 1 was obtained and characterised further. The clone derived from the cipargamin selection performed with Dd2-Polδ parasites (‘Dd2-Polδ-PfATP4^G358S^’) was highly resistant to cipargamin (**Fig. 2**), with an IC_50_ value 1082 ± 138 fold (mean ± SEM, n = 7) higher than that of its Dd2-Polδ parent (**Table 1**).

#### Selections with other PfATP4 inhibitors

We also used Dd2-Polδ parasites to select for parasites resistant to additional PfATP4-associated chemotypes, in each case performing single-step selections with a concentration equating to 6-15 × the IC_50_ for the compound. We exposed Dd2-Polδ parasites (two independent cultures, each containing 2 × 10^8^ parasites) to 250 nM (+)-SJ733. After 16 days, viable parasites were observed in one of the cultures (‘(+)-SJ733-selected Culture 1’; the other culture had no viable parasites on Day 25 and was discarded). We also attempted to generate high-level resistance to two compounds from the Medicines for Malaria Venture’s (MMV’s) ‘Malaria Box’ − MMV665949 and MMV006656. These compounds are structurally unrelated to the spiroindolones, dihydroisoquinolones, pyrazoleamides, or each other, but display the hallmarks of PfATP4 inhibition: an increase in [Na^+^]_cyt_ and pH_cyt_ (24) and inhibition of Na^+^-ATPase activity in parasite membrane preparations (15). We exposed Dd2-Polδ parasites to a single high concentration of 50 μM MMV665949 or 5 μM MMV006656 (two independent cultures, each containing 2 × 10^8^ parasites, for each compound). For MMV665949, resistant parasites emerged from one of the cultures (‘MMV665949-selected Culture 1’) on Day 17. No parasites were observed in the second MMV665949-containing culture within 25 days and the culture was discarded. For MMV006656, no viable parasites were observed and the cultures were discarded on Day 42.

### High-level cipargamin resistance is conferred by a G358S mutation in PfATP4

#### Cipargamin-selected HCR parasites

We extracted genomic DNA from the Dd2-PfATP4^T418N,P990R^ cultures selected with cipargamin (cultures A and B) and from four clonal lines (HCR1-4) obtained from culture B. Sequencing the *pfatp4* gene revealed the same set of mutations in each case (relative to the Dd2 sequence): those encoding the T418N and P990R mutations present in the Dd2-PfATP4^T418N,P990R^ parental line and a third mutation (G1072A) that codes for a G358S mutation in the PfATP4 protein.

We subjected Dd2, Dd2-PfATP4^T418N,P990R^, HCR1 and HCR2 to whole-genome sequencing. This revealed a duplication in chromosome 12 in HCR1 and HCR2 that was not present in Dd2 or Dd2-PfATP4^T418N,P990R^, in a region (12:520kb – 556kb) covering nine genes including *pfatp4*. The *pfatp4* mutation coding for G358S was detected at a frequency of ∼ 50% in both HCR1 and HCR2 (**Fig. S1**), suggesting that the duplication of *pfatp4* occurred before the mutation, and that the mutation is only present in one of the two copies of the gene. Dd2-PfATP4^T418N,P990R^, HCR1 and HCR2 were all confirmed to have the mutations coding for T418N and P990R in PfATP4. Thus, the HCR1 and HCR2 parasites have two copies of *pfatp4*, one encoding the PfATP4^T418N,P990R^ variant present in the parental line and the other encoding the triple-mutant variant PfATP4^G358S,T418N,P990R^. Four other SNPs in different genes, which we considered unlikely to contribute to cipargamin resistance, were also noted in HCR1 and/or HCR2 (**Data S1**). A reduction in the degree of amplification of a region on chromosome 5 containing *pfmdr1* was also observed in HCR2 (**Data S1, Fig. S1**).

#### Cipargamin and (+)-SJ733 selected hypermutator parasites

We also extracted genomic DNA from the Dd2-Polδ bulk cultures selected with cipargamin (cipargamin-selected Culture 1) and (+)-SJ733 ((+)-SJ733-selected Culture 1), and from a clone derived from cipargamin-selected Culture 1, and sequenced the *pfatp4* gene. In all cases we observed a single mutation in the *pfatp4* gene that was not present in the Dd2-Polδ parents – a G1072A mutation in the gene, which codes for the G358S mutation in the PfATP4 protein.

### Response of parasites harbouring the PfATP4 G358S mutation to a variety of antiplasmodial agents

#### Cipargamin-selected HCR parasites

In addition to being highly resistant to cipargamin, the HCR1 and HCR2 parasites (harbouring PfATP4^T418N,P990R^ and PfATP4^G358S,T418N,P990R^) were highly resistant to the dihydroisoquinolone (+)-SJ733 and the pyrazoleamide PA21A050, with the IC_50_ values for the HCR parasites being 54 - 66 fold and 22 - 23 fold higher than those of their Dd2-PfATP4^T418N,P990R^ parents, respectively, and 300 - 366 fold and 50 - 51 fold higher than those of Dd2, respectively (**Table 1**). HCR1 and HCR2 parasites were slightly but significantly *more sensitive* to dihydroartemisinin (the compound formed *in vivo* from all clinically used artemisinin derivatives) than parental and Dd2 parasites (**Table 1**). HCR2 parasites were slightly more resistant to chloroquine than the other parasites (**Table 1**).

#### Cipargamin-selected hypermutator parasites

In addition to being highly resistant to cipargamin, the clone derived from the cipargamin selection performed with Dd2-Polδ parasites (‘Dd2-Polδ-PfATP4^G358S^’) was highly resistant to (+)-SJ733, with an IC_50_ value 237 ± 51 fold (mean ± SEM, n = 4) higher than that of the Dd2-Polδ parent (**Table 1**). Dd2-Polδ-PfATP4^G358S^ parasites were moderately resistant to PA21A050 and MMV006656, with IC_50_ values 5.3 ± 0.1 fold (mean ± SEM, n = 4) and 5.7 ± 1.3 fold (mean ± SEM, n = 4) higher than those of the Dd2-Polδ parent, respectively (**Table 1**). There was no significant change in their susceptibility to MMV665949, chloroquine or dihydroartemisinin (**Table 1**). Thus, the G358S mutation in PfATP4 was associated with a decrease in parasite susceptibility to four out of five compounds for which there is evidence for PfATP4 inhibition, and did not affect parasite susceptibility to the unrelated drugs chloroquine or dihydroartemisinin.

#### Parasites selected with MMV665949

The MMV665949-selected parasites (MMV665949-selected Culture 1; not cloned) were tested for their sensitivity to growth inhibition by MMV665949 and cipargamin. In paired experiments, the IC_50_ value for MMV665949 was 11.0 ± 1.6 fold higher for MMV665949-selected Culture 1 (IC_50_ = 44 ± 7 μM) than for the parental Dd2-Polδ parasites (IC_50_ = 4.3 ± 0.7 μM) (mean ± SEM, n = 4). MMV665949-selected Culture 1 showed only a low level of resistance (2.5 ± 0.5 fold) to cipargamin, with an IC_50_ value of 1.42 ± 0.30 nM, compared to 0.57 ±0.02 nM in paired experiments with the parental Dd2-Polδ parasites (mean ± SEM, n = 3). Thus, while the single-step selections with cipargamin and (+)-SJ733 both resulted in parasites with the G358S mutation in PfATP4 that were highly resistant to both compounds, the single-step selection with the structurally unrelated PfATP4-associated compound MMV665949 did not. The MMV665949-selected parasites were not characterised further in this study.

### Mechanistic basis for high-level resistance to cipargamin caused by the G358S mutation in PfATP4

#### Effect of the G358S mutation in PfATP4 and the equivalent mutation in TgATP4 on the disruption of Na^+^ regulation by cipargamin and (+)-SJ733

PfATP4 is required for the maintenance of a low [Na^+^]_cyt_ in *P. falciparum* parasites, with inhibition of the protein resulting in an increase in [Na^+^]_cyt_ (11). We investigated whether the G358S mutation in PfATP4 rendered parasites resistant to having their [Na^+^]_cyt_ dysregulated by cipargamin and (+)-SJ733. We did this by loading Dd2-Polδ-PfATP4^G358S^ and parental Dd2-Polδ parasites with the Na^+^-sensitive fluorescent dye SBFI, and measuring [Na^+^]_cyt_ in parasites exposed to a range of concentrations of cipargamin and (+)-SJ733. We found that Dd2-Polδ-PfATP4^G358S^ parasites were highly resistant to having their [Na^+^]_cyt_ perturbed by both cipargamin and (+)-SJ733 (**Fig. 3A**,**B**). It was not possible to determine precise IC_50_ values for cipargamin- and (+)-SJ733-mediated Na^+^ dysregulation in Dd2-Polδ-PfATP4^G358S^ parasites due to the very high concentrations that would have been required to obtain a maximal effect. Nevertheless, it is estimated that the IC_50_ values for [Na^+^]_cyt_ dysregulation would be > 230-fold higher for cipargamin, and > 180-fold higher for (+)-SJ733, in Dd2-Polδ-PfATP4^G358S^ parasites than in the parental Dd2-Polδ parasites (**Table 2**).

**Table 2.** Susceptibility of Dd2-Polδ and Dd2-Polδ-PfATP4^G358S^ *P. falciparum* parasites and TgATP4^WT^ and TgATP4^G419S^-HA *T. gondii* parasites to Na^+^ dysregulation and inhibition of ATP4-associated ATPase activity by cipargamin and (+)-SJ733. The IC_50_ values (mean ± SEM) are shown, with the number of independent experiments shown in brackets. For each compound, assay type and parasite species, the IC_50_ values (where possible to obtain) for the ATP4 mutant and its WT counterpart were compared using unpaired t-tests.

|  | IC<sub>50</sub> for Na<sup>+</sup> dysregulation (nM) |  | IC<sub>50</sub> for ATP4-associated ATPase activity (nM) |  |
| --- | --- | --- | --- | --- |
|  | Cipargamin | (+)-SJ733 | Cipargamin | (+)-SJ733 |
| Dd2-Polδ | 8.4 ± 3.0 (5) | 55 ± 15 (4) | 2.5 ± 0.8 (4) | 5.2 ± 2.3 (3) |
| Dd2-Polδ-PfATP4<sup>G358S</sup> | > 2000 (6) | > 10,000 (4) | 99 ± 21 (4) (** *P* = 0.004) | 829 ± 25 (3) (*** *P* < 0.001) |
| WT *T. gondii* (TgATP4<sup>WT</sup>) | 110 ± 3 (3) | 5045 ± 572 (4) | 8.4 ± 5.8 (4) | 17.8 ± 9.2 (3) |
| TgATP4<sup>G419S</sup>-HA | > 8000 (3) | > 40,000 (3) | 288 ± 111 (4) (* *P* = 0.05) | > 4000 (3) |

**Fig. 3.**
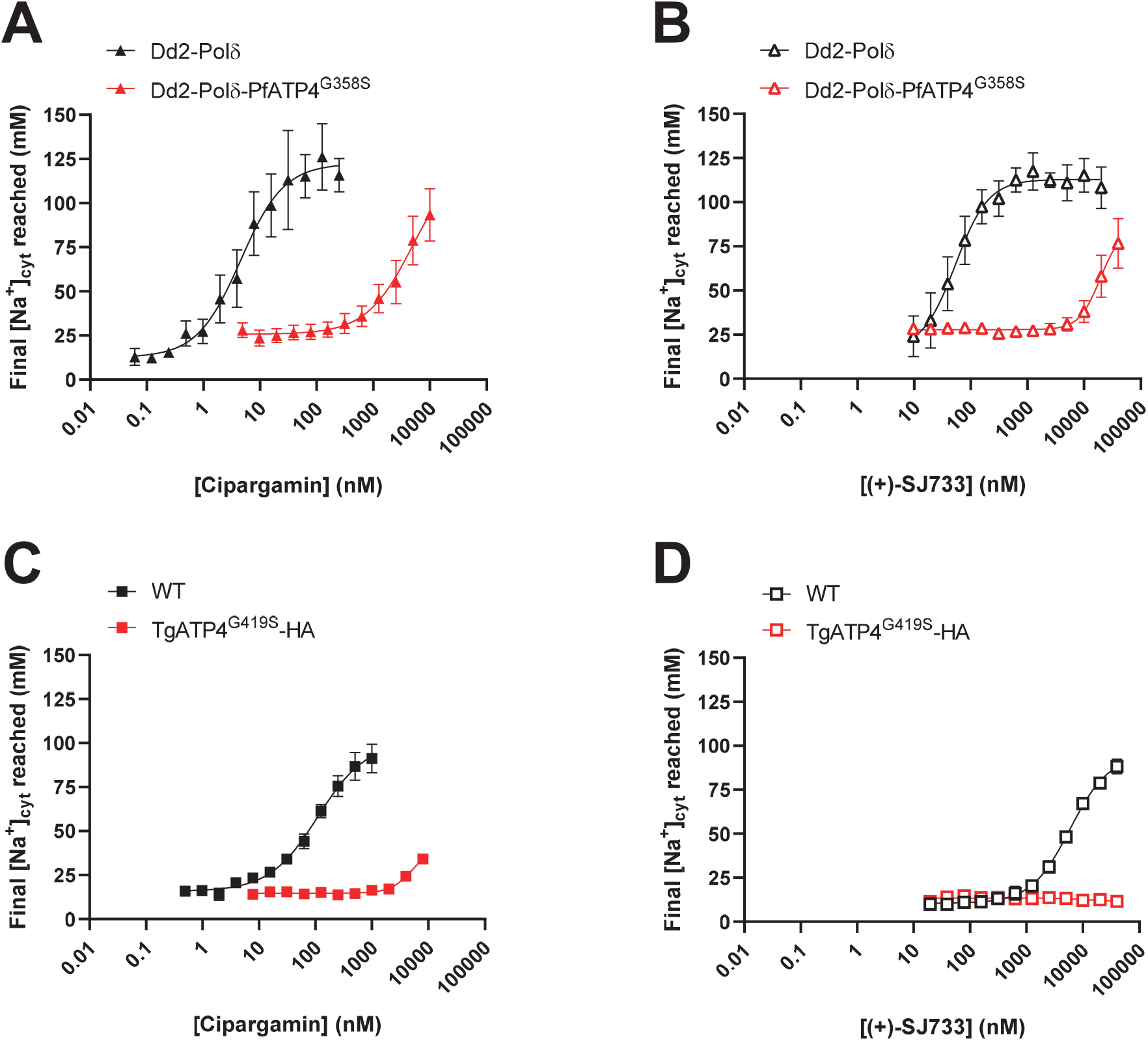
Parasites with the G358S mutation in PfATP4 or the G419S mutation in TgATP4 are resistant to cipargamin- and (+)-SJ733-mediated Na^+^ dysregulation. The final [Na^+^]_cyt_ reached after a ∼ 90 min exposure to a range of concentrations of cipargamin (**A** and **C**) or (+)-SJ733 (**B** and **D**). The measurements were performed at 37°C in pH 7.1 Physiological Saline Solution with isolated SBFI-loaded Dd2-Polδ (black) and Dd2-Polδ-PfATP4^G358S^ (red) *P. falciparum* trophozoites (**A** and **B**) and with extracellular SBFI-loaded *T. gondii* tachyzoites (**C** and **D**) expressing TgATP4^WT^ (black) or TgATP4^G419S^-HA (red). The data are the mean (± SEM) from the following number of independent experiments: 3-6 (**A**), 3-4 (**B**), 3 (**C**) and 3-4 (**D**). Where not shown, error bars fall within the symbols.

We also investigated the effects of cipargamin and (+)-SJ733 on Na^+^ regulation in the related apicomplexan parasite *T. gondii*. We used CRISPR-Cas9 to generate *T. gondii* parasites (‘TgATP4^G419S^-HA’) that harbour the equivalent mutation – G419S – in TgATP4 (**Fig. S2**). We isolated successfully-edited clones (**Fig. S3**) and confirmed that the parasites expressed TgATP4^G419S^-HA at the plasma membrane (**Fig. S4**). In contrast to *P. falciparum, T. gondii* does not require TgATP4 activity for growth (18), and the much higher concentrations of cipargamin needed to kill *T. gondii* parasites (38) likely do so via off target effects. However, TgATP4 is required for the maintenance of a low [Na^+^]_cyt_ in extracellular tachyzoites under physiological (∼ 130 mM extracellular Na^+^) conditions (18). We investigated the potency with which cipargamin and (+)-SJ733 dysregulate [Na^+^]_cyt_ in extracellular *T. gondii* tachyzoites (TATi/*Δku80* strain) expressing wild-type TgATP4 (TgATP4^WT^) and TgATP4^G419S^-HA. These assays revealed that TgATP4^G419S^-HA parasites were much less sensitive to having their [Na^+^]_cyt_ dysregulated by cipargamin and (+)-SJ733 than TgATP4^WT^ parasites (**Fig. 3C**,**D**).

#### Effect of the G358S mutation in PfATP4 and the equivalent mutation in TgATP4 on inhibition of ATP4 ATPase activity by cipargamin and (+)-SJ733

To date, the most direct assay available for measuring PfATP4 activity entails measuring Na^+^-dependent ATPase activity in *P. falciparum* membrane preparations (11,15). Approximately 25% of the total ATPase activity measured in *P. falciparum* membrane preparations is Na^+^-dependent. Through studies with PfATP4 inhibitors and *pfatp4* mutant lines, this Na^+^-dependent ATPase activity has been attributed to PfATP4 (11,15). We adapted the ATPase assay for use with *T. gondii* and detected a cipargamin-sensitive Na^+^-dependent ATPase activity in *T. gondii* membrane preparations (**Fig. 4C**,**D**).

**Fig. 4.**
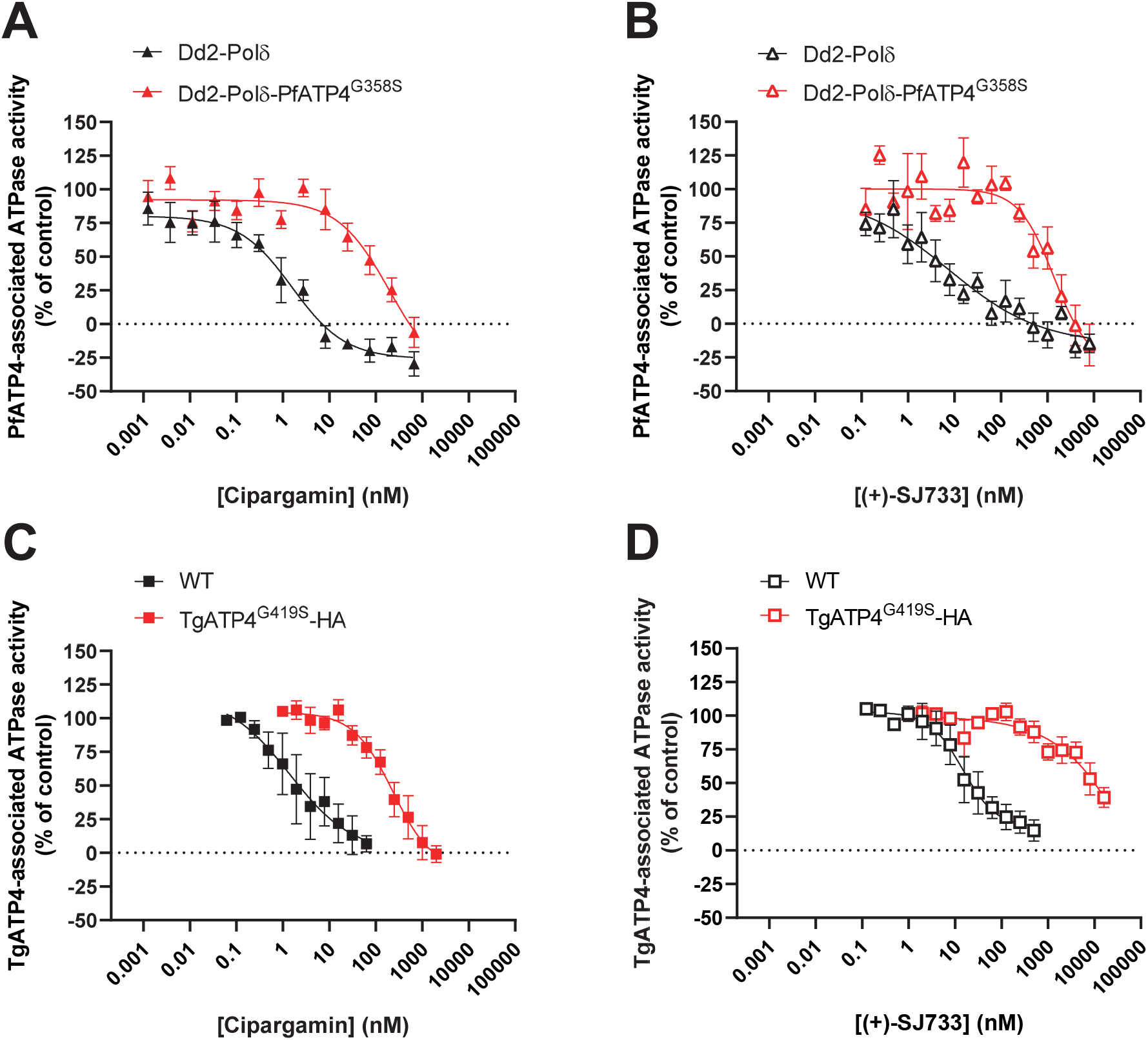
Reduced sensitivity of PfATP4^G358S^- and TgATP4^G419S^-associated ATPase activity to inhibition by cipargamin and (+)-SJ733. The potency by which cipargamin (**A** and **C**) and (+)-SJ733 (**B** and **D**) inhibit ATP4-associated ATPase activity was determined using membranes prepared from Dd2-Polδ (black) and Dd2-Polδ-PfATP4^G358S^ (red) *P. falciparum* trophozoites (**A** and **B**) or *T. gondii* tachyzoites (**C** and **D**) expressing TgATP4^WT^ (black) or TgATP4^G419S^-HA (red). The data are the mean (± SEM) from the following number of independent experiments (each performed on different days with different membrane preparations): 4 (**A**), 3 (**B**), 4 (**C**) and 3 (**D**). Where not shown, error bars fall within the symbols.

We investigated whether the G358S mutation in PfATP4, and the G419S mutation in TgATP4, rendered ATP4-associated ATPase activity less sensitive to inhibition by cipargamin and (+)-SJ733. Using membranes prepared from Dd2-Polδ-PfATP4^G358S^ and their parental Dd2-Polδ parasites, we found that PfATP4^G358S^-associated ATPase activity was 40-fold and 159-fold less sensitive to inhibition by cipargamin and (+)-SJ733, respectively, than PfATP4^WT^-associated ATPase activity (**Fig. 4A**,**B; Table 2**). A similar result was obtained with membranes prepared from TgATP4^G419S^-HA parasites and WT parasites, with TgATP4^G419S^-HA-associated ATPase activity displaying a 34-fold and > 225-fold lower susceptibility to cipargamin and (+)-SJ733, respectively, than TgATP4^WT^-associated ATPase activity (**Fig. 4C**,**D; Table 2**). These results provide strong evidence that the G358S mutation in PfATP4, and the equivalent G419S mutation in TgATP4, greatly reduce the sensitivity of the proteins to inhibition by cipargamin and (+)-SJ733.

### Effect of the G358S mutation on PfATP4 function

From a clinical perspective it is important to ascertain whether the G358S mutation in PfATP4 impairs the protein’s function, and whether or not this leads to a defect in parasite growth. Membrane ATPase assays have been used previously to compare the biochemical properties of Dd2-PfATP4 and Dd2-PfATP4^T418N,P990R^, revealing that the T418N and P990R mutations are associated with a slight decrease in the affinity of PfATP4 for Na^+^ (i.e. an elevated K_m_(Na^+^)), but no change in its maximum rate (V_max_) (15). We investigated the Na^+^-dependence of PfATP4-associated membrane ATPase activity in membranes prepared from Dd2-Polδ and Dd2-Polδ-PfATP4^G358S^ parasites (**Fig. 5A**). With membranes prepared from Dd2-Polδ-PfATP4^G358S^ parasites, we estimated a K_m_(Na^+^) for PfATP4^G358S^ of 75 ± 17 mM (mean ± SEM, n = 5; **Fig. 5A**). This was significantly higher than the K_m_(Na^+^) estimated for WT PfATP4 in the parental Dd2-Polδ parasites, which was 25.8 ± 1.8 mM (mean ± SEM, n = 5; *P* = 0.04, paired t-test; **Fig. 5A**). The V_max_ values estimated for WT PfATP4 and PfATP4^G358S^ were not significantly different from one another (31.7 ± 3.0 and 30.5 ± 3.9 nmol P_i_ per mg of (total) protein per min, respectively; mean ± SEM, n = 5; *P* = 0.6, paired t-test).

**Fig. 5.**
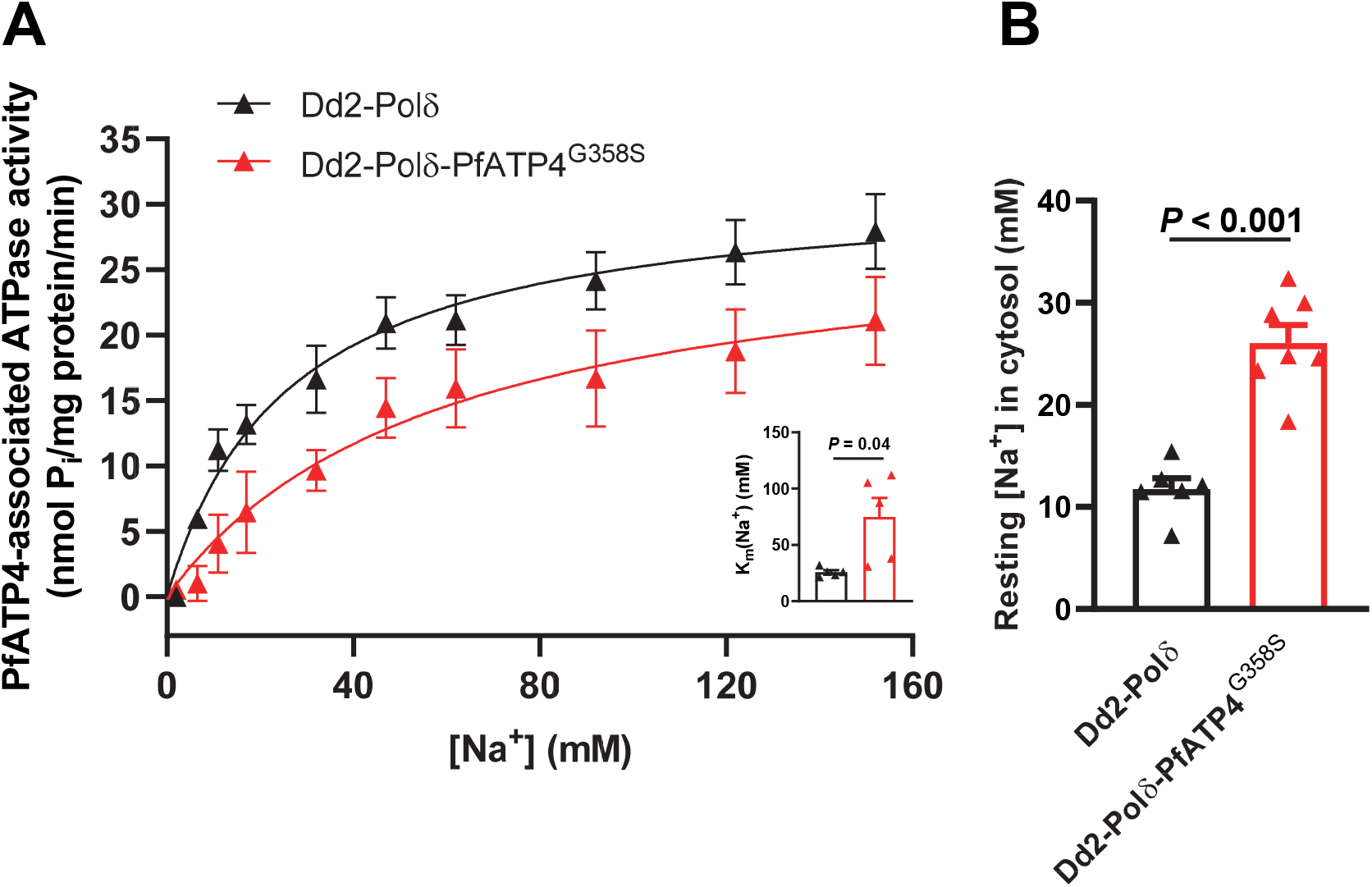
The G358S mutation in PfATP4 affects the Na^+^-dependence of PfATP4-associated ATPase activity. **A**. Effect of [Na^+^] on PfATP4-associated ATPase activity in membranes prepared from Dd2-Polδ-PfATP4^G358S^ parasites (red) and their Dd2-Polδ parents (black). The data shown are the mean (± SEM) from five independent experiments, each performed on different days with different membrane preparations. The Michaelis-Menten equation (PfATP4-associated ATPase activity = V_max_ × [Na^+^]/([Na^+^] + K_m_(Na^+^))) was fitted to the data. The K_m_(Na^+^) in the two lines is shown in the inset, with the symbols showing the K_m_(Na^+^) values from each individual experiment, the bars showing the mean, and the error bars showing the SEM. A paired t-test was used to compare the K_m_(Na^+^) values obtained for the two lines. **B**. Resting cytosolic [Na^+^] in Dd2-Polδ-PfATP4^G358S^ parasites (red) and their Dd2-Polδ parents (black). The measurements were performed with isolated trophozoite-stage parasites loaded with the Na^+^-sensitive dye SBFI, and suspended in pH 7.1 Physiological Saline Solution at 37°C. The bars show the mean + SEM obtained from six or seven independent experiments for each line, and the symbols show the results obtained in individual experiments. The *P* value is from an unpaired t-test.

Consistent with the finding of a reduced affinity of PfATP4^G358S^ for Na^+^, we found that Dd2-Polδ-PfATP4^G358S^ parasites had a significantly (2.2 fold) higher resting [Na^+^]_cyt_ than their Dd2-Polδ parents (**Fig. 5B**). The resting [Na^+^]_cyt_ concentrations for the Dd2-Polδ and Dd2-Polδ-PfATP4^G358S^ parasites were 11.7 ± 1.1 mM (mean ± SEM, n = 6) and 26.0 ± 1.8 mM (mean ± SEM, n = 7), respectively.

Of the six PfATP4-mutant parasite lines for which resting [Na^+^]_cyt_ values have been reported previously, five (PfATP4^A211T^, PfATP4^I203L/P990R^, PfATP4^L350H^, PfATP4^I398F/P990R^ and PfATP4^T418N,P990R^) were found to have an elevated resting [Na^+^]_cyt_ compared to their parents (11,20,23). We confirmed that the Dd2-PfATP4^T418N,P990R^ parent of the HCR clones had a higher [Na^+^]_cyt_ than its Dd2 parent (**Fig. S5**). For HCR1 and HCR2 parasites, the mean [Na^+^]_cyt_ was slightly higher than that of their Dd2-PfATP4^T418N,P990R^ parent, but this was not statistically significant (**Fig. S5**).

To investigate further how frequently mutations in PfATP4 are associated with an elevation of the parasite’s resting [Na^+^]_cyt_ we tested an additional five PfATP4-mutant lines that were selected previously with various PfATP4 inhibitors (PfATP4^T418N^, PfATP4^Q172K^, PfATP4^A353E^, PfATP4^P966S^ and PfATP4^P966T^). Four of the five lines had a significantly (*P* < 0.05) higher resting [Na^+^]_cyt_ than their parents (**Fig. S5**). Thus, considering all the lines expressing a single mutant variant of *pfatp4* for which resting [Na^+^]_cyt_ measurements have now been performed, 10 of the 12 mutant variants of PfATP4, including PfATP4^G358S^, have been associated with an elevation of resting [Na^+^]_cyt_.

### Effect of the G358S mutation in PfATP4 on the growth of asexual blood stage parasites

To determine whether the G358S mutation in PfATP4 was associated with a cost to the fitness of asexual blood stage parasites, we investigated the *in vitro* growth of Dd2-Polδ-PfATP4^G358S^ parasites and their Dd2-Polδ parents in the absence of any drug pressure (**Fig. 6**). The ∼ 48 h *P. falciparum* erythrocytic cycle begins when a ‘merozoite’ invades an erythrocyte. The intraerythrocytic merozoite then develops into a ring-stage parasite, and increases in volume as it matures into a young trophozoite, mature trophozoite and then a schizont. The schizont segments to form daughter parasites, which egress from the erythrocyte and commence the next cyle. In general, a change in growth rate can result from either a decrease in the number of viable parasites produced per cycle or from a lengthening of the cycle. To test for these possibilities, we: (1) adjusted parasite cultures to a parasitaemia of 1% and measured the parasitaemias after one complete cycle; and (2) measured the change in volume distribution of trophozoites after intervals of ∼ 48 h.

**Fig. 6.**
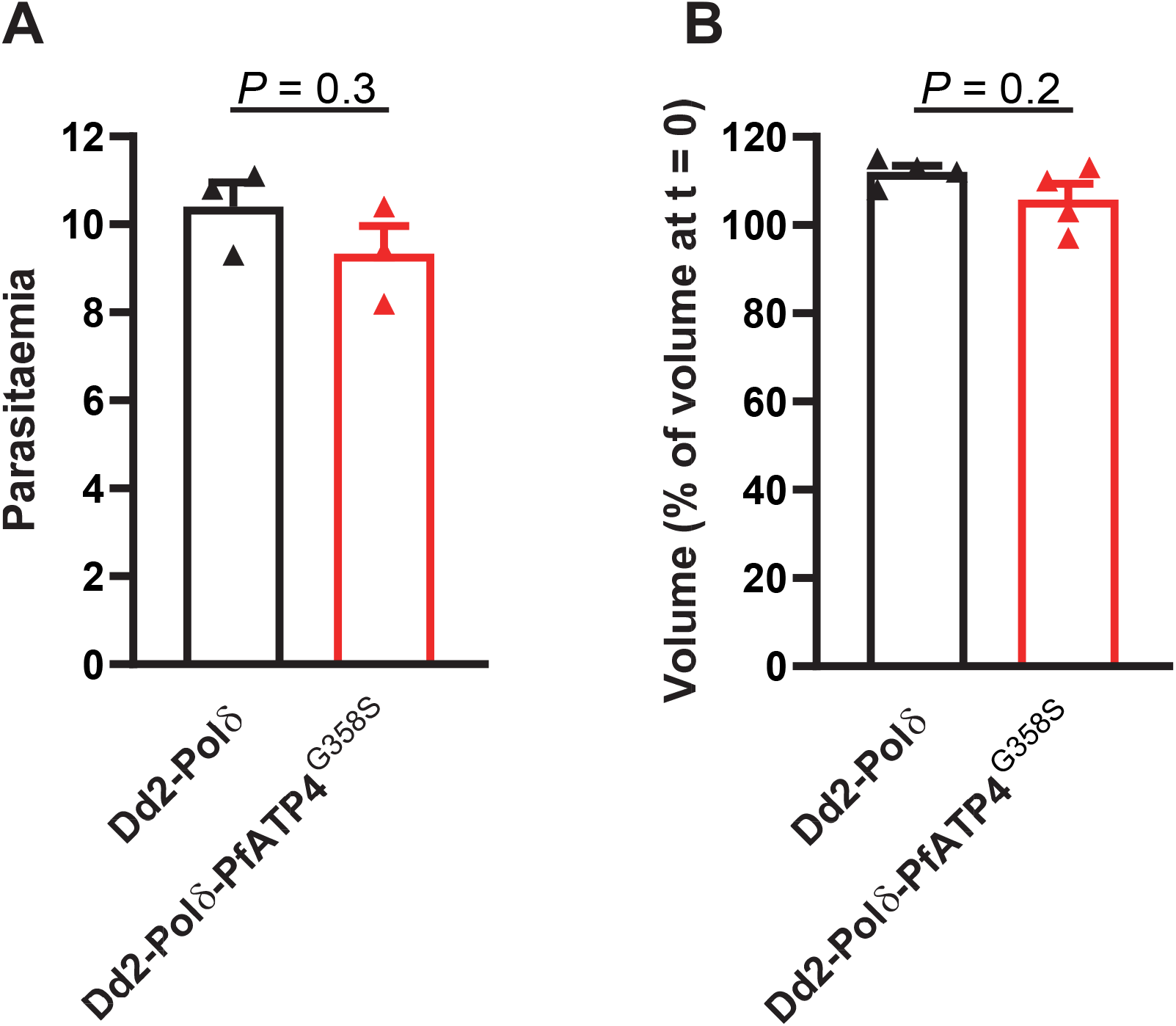
Production of viable daughter parasites and progression through the erythrocytic cycle is similar in Dd2-Polδ-PfATP4^G358S^ and Dd2-Polδ parasites. A. The parasitaemias obtained one cycle (∼ 48 h) after adjusting the parasitaemia of cultures to 1%. The bars show the mean from three independent experiments, each performed with three cultures (technical replicates). **B**. Parasite volume (as a percentage of that measured at 0 h) after one cycle (48 h) of growth. The bars show the mean from four independent experiments, each performed with at least two technical replicates. In **A** and **B**, the error bars are SEM, and the symbols show the data from each independent experiment. The *P* values are from unpaired t-tests.

To compare the number of viable parasites produced per cycle by Dd2-Polδ-PfATP4^G358S^ parasites and their Dd2-Polδ parents, we used flow cytometry to determine the parasitaemia in three replicate cultures containing synchronous trophozoite-stage parasites, then diluted each one to a parasitaemia of 1%. The parasitaemias in each culture were then determined approximately 48 h later using flow cytometry. This was performed on three independent occasions, yielding the data shown in **Fig. 6A**. One cycle (∼ 48 h) after being adjusted to a parasitaemia of 1%, the parasitaemia reached 10.4 ± 0.6% in the Dd2-Polδ cultures, and 9.3 ± 0.6% in the Dd2-Polδ-PfATP4^G358S^ cultures (mean ± SEM, n = 3; *P* = 0.3, unpaired t-test), consistent with Dd2-Polδ and Dd2-Polδ-PfATP4^G358S^ parasites producing a similar number of viable daughter parasites per cycle. This is consistent with our observation that there was no obvious difference in the growth of these parasites during the routine culture of the lines.

To compare the rate of progression through the asexual life cycle in Dd2-Polδ and Dd2-Polδ-PfATP4^G358S^ parasites, we measured the mean volume of isolated trophozoite-stage parasites, then determined the change in mean volume 48 h later (**Fig. 6B**). As a percentage of the starting volume, the mean trophozoite volume after 48 h was 112 ± 1% for Dd2-Polδ parasites, and 106 ± 4% for the Dd2-Polδ-PfATP4^G358S^ parasites (mean ± SEM, n = 4; *P* = 0.2, unpaired t-test). The small increase in mean parasite volume between the 0 h and 48 h time points observed for both lines suggests that the duration of their erythrocytic life cycles is slightly shorter than 48 h.

Thus, despite affecting the function of PfATP4, the G358S mutation was not associated with a significant change in the length of the erythrocytic cycle or in the number of viable merozoites produced per cycle during the asexual blood stage.

## Discussion

PfATP4 has emerged as a major antimalarial drug target, with the PfATP4 inhibitor cipargamin having undergone extensive testing in Phase 1 and Phase 2 clinical trials (27), and another PfATP4 inhibitor (+)-SJ733 also having been tested in humans (26). Understanding the likelihood and mechanisms of resistance to PfATP4 inhibitors is important as compounds with this mechanism are pursued further. In this study we generated highly cipargamin-resistant parasites in three independent selections with cipargamin: two stepwise selections commencing with low-level resistant parasites, and one single-step selection commencing with parasites displaying a hypermutator phenotype. In each case these selections gave rise to parasites with a G358S mutation in PfATP4. A fourth independent experiment performed with Dd2-Polδ in a different laboratory also yielded the same result (Kümpornsin *et al*., in preparation). In contrast, many different mutations in PfATP4 have been reported in parasites selected under conditions in which the acquisition of a low level of resistance to cipargamin is sufficient for survival (9,34). Thus, it would appear that there are many roads to low-level cipargamin resistance but few roads to high-level cipargamin resistance. The frequency of resistance, measured at ∼ 2 × 10^−8^ to 1 × 10^−7^ for low-level cipargamin resistance (34), decreased to < 6.6 × 10^−9^ for high-level resistance (in Dd2-PfATP4^T418N,P990R^). Indeed, we only succeeded in generating highly resistant parasites via a single-step procedure using genetically engineered parasites with mutations in DNA polymerase δ and a hypermutator phenotype. Based on studies with *P. berghei* parasites similarly engineered to have mutations in DNA polymerase δ (39), the Dd2-Polδ parasites might be expected to have a mutation rate ∼ 80-fold higher than those of wild-type Dd2 parasites.

In a recent study, the G358S mutation in PfATP4 was observed in recrudescent parasites that emerged in a clinical trial for cipargamin (40). The G358S mutation was observed in 22/25 instances of recrudescence. On five of these occasions there was a mix of G358S with WT sequence at the positions tested, G358A or G359A. There were also three instances of recrudescence in which the G358S mutation was not observed. These were associated (on one occasion each) with the G359A, L354V and L181F mutations in PfATP4 (40). The results of this trial demonstrate the clinical significance of the G358S mutation and provide further evidence that it may be one of few mutations that can confer high-level resistance to cipargamin. It will be interesting to investigate the G358A, G359A, L354V and L181F mutations in future studies to assess the level of resistance conferred by those mutations and their effects on PfATP4 function and parasite fitness.

In contrast to cipargamin, for which high-level resistance had not been observed previously in *in vitro* evolution studies, high-level resistance to SJ733 has been encountered before and there appear to be multiple mutations that can confer this phenotype (23,41). Genetically engineered parasites bearing the L350H or P412T mutations in PfATP4 have been shown to display a high level of resistance to SJ733, with IC_50_s of 3.5 μM and 10 μM, respectively, compared to 75 nM for parental wild-type parasites (41). With IC_50_s > 17 μM for (+)-SJ733, the HCR parasites generated in this study have the highest (+)-SJ733 IC_50_s reported to date. The G358S mutation in PfATP4 has been reported once before in parasites selected with SJ733 (23). Few details have been published for these parasites, although they were reported to display only a 5-fold resistance to SJ733 (23), which is much lower than the (237 ± 51 fold; mean ± SEM, n = 4) difference in (+)-SJ733 sensitivity observed between Dd2-Polδ and Dd2-Polδ-PfATP4^G358S^ in this study.

The HCR parasites generated in this study also have the highest level of PA21A050 resistance reported to date (mean IC_50_s of ∼ 60 nM). Previous selections with a pyrazoleamide yielded parasites with an IC_50_ for PA21A050 of 16 nM (compared to 0.7 nM for the parental line) and with mutations in five proteins, including PfATP4 (V178I) and the Ca^2+^-dependent protein kinase PfCDPK5 (T392A) (25). Whole-genome sequencing of our HCR parasites did not reveal any mutations in the gene encoding PfCDPK5 (or in the three other genes that were mutated along with PfATP4 in the pyrazoleamide-selected lines) relative to their parents.

The HCR parasites were more resistant to cipargamin, (+)-SJ733 and PA21A050 than the Dd2-Polδ-PfATP4^G358S^ parasites. The two mutations in Dd2-PfATP4^T418N,P990R^ parasites conferred some resistance to each drug (ranging from 2-14-fold), with the fold difference in IC_50_ value between HCR parasites and their Dd2 ‘grandparents’ equating to approximately the product of the fold difference between Dd2 and Dd2-PfATP4^T418N,P990R^ parasites (resulting from the T418N and P990R mutations in PfATP4) and that between Dd2-PfATP4^T418N,P990R^ and HCR parasites (stemming from the amplification of *pfatp4* and the presence of the G358S-coding mutation in one of the copies). The G358S mutation in PfATP4 was an important contributor to high-level resistance for cipargamin, with Dd2-Polδ-PfATP4^G358S^ parasites having a 1081 ± 138 fold higher IC_50_ (mean ± SEM, n = 7) than their parents (c.f. a ∼ 4200-fold difference between HCR parasites and Dd2). For (+)-SJ733, the G358S mutation was also a major contributor to high-level resistance, with Dd2-Polδ-PfATP4^G358S^ parasites having a 237 ± 51 fold higher IC_50_ (mean ± SEM, n = 4) than their parents (c.f. a 300-366 fold difference between HCR parasites and Dd2). The contribution of the G358S mutation was less pronounced for PA21A050, with the Dd2-Polδ-PfATP4^G358S^ parasites displaying only a 5.3 ± 0.1 fold increase in IC_50_ (mean ± SEM, n = 4) (c.f. a ∼ 50-fold difference between HCR parasites and Dd2). The T418N and P990R mutations in PfATP4 were associated with a 2.2 ± 0.2 fold increase in PA21A050 IC_50_ (mean ± SEM, n = 3). Perhaps the amplification of *pfatp4* played a proportionally greater role in resistance to PA21A050 than in resistance to cipargamin or (+)-SJ733.

Relative to their Dd2-PfATP4^T418N,P990R^ parent, HCR1 and HCR2 parasites were both significantly more sensitive to dihydroartemisinin (both with ∼ 1.6-fold lower IC_50_s), and HCR2 was significantly less susceptible to chloroquine (1.3-fold higher IC_50_). The reasons for these differences are not clear, although for HCR2 parasites, the reduction in *pfmdr1* copy number relative to HCR1 and Dd2-PfATP4^T418N,P990R^ parasites (**Fig. S1**) might play a role (42). In any case, the finding that there was no difference in the sensitivity of Dd2-Polδ and Dd2-Polδ-PfATP4^G358S^ parasites to chloroquine or dihydroartemisinin suggests that the G358S mutation in PfATP4 is not responsible.

Our investigations into the mechanism of resistance revealed that the G358S mutation in PfATP4, and the G419S mutation in TgATP4, rendered the Na^+^-ATPase activity of the proteins resistant to inhibition by cipargamin and (+)-SJ733. Comparing the IC_50_s for Na^+^-ATPase inhibition, PfATP4^G358S^ was 40-fold less sensitive to inhibition by cipargamin than PfATP4^WT^, and TgATP4^G419S^-HA was 34-fold less less sensitive than TgATP4^WT^. For (+)-SJ733 the fold differences were 159 and > 225, respectively. TgATP4^WT^-associated ATPase activity was about 3-fold less sensitive to inhibition by both cipargamin and (+)-SJ733 than PfATP4^WT^-associated ATPase activity. (+)-SJ733 was only ∼ 2-fold less potent than cipargamin against both PfATP4^WT^ and TgATP4^WT^ (as measured in cell-free assays), yet the difference in potency between the drugs was greater (> 6-fold) in [Na^+^]_cyt_ assays (performed with intact isolated *P. falciparum* or extracellular *T. gondii* parasites) and in parasite proliferation assays with *P. falciparum* parasites. This raises the possibility that the relative intracellular concentration reached for (+)-SJ733 is lower than that for cipargamin.

*T. gondii* proved useful as a model with which to study ATP4 activity in this study. There is significant homology between TgATP4 and PfATP4, which allowed the residue equivalent to G358 in PfATP4 to be identified unequivocally in TgATP4. Furthermore, the recent characterisation of TgATP4 provided evidence that the protein performs the same role that has been ascribed to PfATP4 (18). *T. gondii* is highly genetically tractable, with clonal TgATP4^G419S^-HA parasites generated within weeks. Furthermore, *T. gondii* parasites can survive and proliferate in the absence of TgATP4 activity, which can be exploited to study mutations in ATP4 that impair its function. Thus, *T. gondii* might serve as a good model for the investigation of a variety of other resistance-conferring mutations in ATP4, as well as to probe other residues that might be important for ATP4 function.

The G358S mutation in PfATP4 affected the function of the protein. Studies of PfATP4-associated ATPase activity revealed that the affinity of PfATP4^G358S^ for Na^+^ was lower than that of PfATP4^WT^. The increase in K_m_(Na^+^) (relative to (Dd2) PfATP4^WT^) was ∼ 3-fold, significantly greater than the 1.3-fold elevation in K_m_(Na^+^) seen for the only other PfATP4 mutant that has been investigated in this way, PfATP4^T418N,P990R^ (15). Consistent with the finding of a reduced affinity of PfATP4^G358S^ for Na^+^, Dd2-Polδ-PfATP4^G358S^ parasites had an elevated resting [Na^+^]_cyt_ compared to their parents. For HCR parasites, the resting [Na^+^]_cyt_ appeared slightly increased beyond the elevated [Na^+^]_cyt_ caused by the other two mutations (T418N and P990R) found in these parasites and their Dd2-PfATP4^T418N,P990R^ parents, but this was not significant. In total, in this study and others (11,20,23), 12 parasites having a (single) mutant variant of PfATP4 have now been investigated in Na^+^ assays, with 10 of these found to have a significantly elevated [Na^+^]_cyt_.

We did not observe obvious differences in the growth of Dd2-Polδ-PfATP4^G358S^ parasites and their parents during routine culture, and when seeded at 1% parasitaemia, both lines produced a similar increase in parasitaemia after one cycle of growth. There was no clear difference in their rate of progression through the asexual intraerythrocytic cycle, as determined by measuring parasite volume at 48 h intervals. Among other PfATP4 mutant lines, two have been reported to have a growth defect, and three have been reported not to (20,23). From this study and previous studies (20,23), there are now six PfATP4 mutant lines for which both resting [Na^+^]_cyt_ data and growth data are available. Of the four that were found to have normal growth, three (G358S, A211T, I203L/P990R) were found to have an elevated resting [Na^+^]_cyt_ and one (A187V) was found not to. Of two lines reported to have a growth defect, one (L350H) was found to have a ∼ 2.8-fold elevated resting [Na^+^]_cyt_ (23) and one (P996T) was found to have a slight (∼ 1.3 fold) elevation of [Na^+^]_cyt_ that was not statistically significant (**Fig. S5**). Thus, an elevation in resting [Na^+^]_cyt_ does not consistently give rise to a defect in parasite growth, at least during the asexual blood stage.

In conclusion, it is possible for parasites to acquire a clinically significant level of resistance to both cipargamin and (+)-SJ733 while maintaining a robust growth rate, and it will be important to monitor *pfatp4* codon 358 closely in the field. Parasites with the G358S mutation in PfATP4 remained fully susceptible to other antimalarials with unrelated modes of action (chloroquine and dihydroartemisinin). For one PfATP4-inhibiting chemotype, MMV006656, our attempt to select for highly resistant parasites with a 15× IC_50_ concentration of the compound was not successful even with Dd2-Polδ parasites, raising the possibility that there are some PfATP4-inhibiting chemotypes for which parasites may not be able to acquire high-level resistance. Furthemore, the presence of the single G358S mutation had very different effects for different anti-PfATP4 compounds, ranging from a >1000-fold decrease in potency (cipargamin), to no significant change in potency (MMV665949). Thus, in prioritising PfATP4 inhibitors for clinical development in the future, it would be useful to attempt to generate high-level resistance to them with Dd2-Polδ parasites, as well as test them against parasites with the G358S mutation in PfATP4.

## Methods

### *P. falciparum* culture

*P. falciparum* parasites were cultured in human erythrocytes (43), with continuous shaking (44), and were synchronised by sorbitol treatment (45). The cultures, which typically had a haematocrit between 2-4%, were maintained at 37°C in a low-O_2_ atmosphere (1% O_2_, 3% CO_2_ and 96% N_2_). The culture medium was RPMI 1640 containing 25 mM HEPES (Gibco) supplemented with 11 mM additional glucose, 0.2 mM hypoxanthine, 20 μg/mL gentamicin sulphate and 3 g/L Albumax II. The use of human blood in this study was approved by the Australian National University Human Research Ethics Committee. The blood was provided by Australian Red Cross Lifeblood without disclosing the identities of the donors.

### *P. falciparum* lines

A variety of different *P. falciparum* lines were used and generated during this study. Dd2-PfATP4^T418N,P990R^ and its Dd2 parent were described previously (9) and kindly provided by E. A. Winzeler. Dd2-PfATP4^T418N,P990R^ was used to generate highly cipargamin resistant parasites (as described in the Results and shown in **Fig. 1**), from which clones HCR1 and HCR2 were derived by a limiting dilution procedure described previously (46). Dd2-Polδ is a parasite line engineered to have a mutant form of DNA polymerase δ. It will be the subject of a separate study (Kümpornsin *et al*., in preparation). Dd2-Polδ was used to generate the Dd2-PfATP4^G358S^ parasites, as well as parasites displaying high-level resistance to (+)-SJ733 and MMV665949 (as described in the Results).

Clonal Dd2-Polδ-PfATP4^G358S^ parasites were obtained using a FACS-based method. Briefly, erythrocytes infected with late trophozoite-stage parasites were enriched using a Miltenyi Biotec VarioMACS magnet. A FACS Aria III system (Imaging & Cytometry Facility, The John Curtin School of Medical Research, Australian National University) was used to add single cells to individual wells in a 96 well plate containing culture medium and uninfected erythrocytes (2% haematocrit). Cultures were provided with fresh medium and erythrocytes as required. After 18 days, wells containing parasites were identified by assaying for parasite-specific lactate dehydrogenase activity (46).

Resting [Na^+^]_cyt_ measurements for additional *P. falciparum* lines are shown in **Fig. S5**. W2-PfATP4^P966S^, W2-PfATP4^P966T^ and their W2 parent were reported previously (23) and generously provided by R. K. Guy. Dd2-PfATP4^Q172K^, Dd2-PfATP4^A353E^, Dd2-PfATP4^T418N^ and their parent (Dd2 clone 10A) were generated through *in vitro* evolution experiments performed previously (24). The Dd2-PfATP4^T418N^ clone and its parent were described previously (15). Clones of Dd2-PfATP4^Q172K^ and Dd2-PfATP4^A353E^ were obtained from MMV011567-pressured cultures 1 and 2 (24) by limiting dilution.

### *P. falciparum* genomic DNA isolation and sequencing

Genomic DNA was isolated from saponin-isolated trophozoite-stage parasites using either a QIAGEN DNeasy Plant Mini Kit or a QIAGEN DNeasy Blood and Tissue Kit. The primers used for *pfatp4* amplification and sequencing are shown in **Table S1**.

Whole genome sequencing was performed as described previously (47). Resulting fastq files were aligned using Bowtie2 (version 2.2.5) (48) with parameters sensitive-local and maxins 1000. Duplicate reads were removed using Picard tools MarkDuplicates (version 2.2.2). Calling of single nucleotide variants (SNVs) and indels was performed with SNVer (v0.5.3) (49) and VarScan (v2.4) (50). Copy number analysis was performed using the R package QDNaseq (version 1.10.0) (51). Structural variant calling was performed using GRIDSS (version 1.5) (52). Paired-end reads were assembled and aligned to two reference genomes, Dd2 (assembly ASM14979v1) and 3D7 (PlasmoDB-29_Pfalciparum3D7). Mappability, coverage and alignment statistics were better with alignment to the 3D7 reference genome, and this was the version used in analysis. Coverage of all samples was equal or greater than 130X (**Data S1**).

### Isolation of *P. falciparum* parasites from their host erythrocytes

Prior to the preparation of parasite membranes or to measurements of [Na^+^]_cyt_ or cell volume, mature trophozoite-stage parasites (approximately 34-40 h post-invasion) were functionally isolated from their host erythrocytes by brief exposure (of cultures at approximately 4% haematocrit) to saponin (0.05% w/v, of which ≥ 10% was the active agent sapogenin) (53). The parasites were then washed several times in bicarbonate-free RPMI 1640 supplemented with 11 mM additional glucose, 0.2 mM hypoxanthine and 25 mM HEPES (pH 7.10), and maintained in this medium at a density of ∼ 1 × 10^7^ – 3 × 10^7^ parasites mL^-1^ at 37°C until immediately before their use in experiments.

### *P. falciparum* proliferation assays

The effects of compounds of interest on parasite proliferation was measured in 96-well plates, essentially as described previously (54), with a fluorescent DNA-intercalating dye used to detect surviving parasites (55). The assays were initiated with erythrocytes infected with predominantly ring-stage parasites, and the starting parasitaemia and haematocrit were both 1%. The duration of the assays was 72 h and the concentration of DMSO did not exceed 0.1% (v/v). IC_50_ values were determined by fitting a sigmoidal curve to the data (SigmaPlot, Systat Software): *y* = (*a*/(1+(*x*/*x*_*0*_)^*b*^, where *y* is ‘% parasite proliferation’, *x* is the concentration of the test compound, *x*_*0*_ is the IC_50_ value, and *a* and *b* are fitted constants.

### *T. gondii* culture

*T. gondii* parasites were cultured in human foreskin fibroblasts in a humidified 37°C incubator containing 5% CO_2_. Infected host cells were cultured in Dulbecco’s modified Eagle’s medium (DMEM) supplemented with 1% v/v foetal calf serum, 50 U/ml penicillin, 50 µg/ml streptomycin, 10 µg/ml gentamicin, 0.25 µg/ml amphotericin B, 0.2 mM L-glutamine, and 2.0 g/L Na_2_HCO_3_.

### Generating *T. gondii* parasites with a G419S mutation in TgATP4

To generate a strain of *T. gondii* parasites wherein the *tgatp4* gene was mutated to express a TgATP4 protein containing the G419S mutation, we used a CRISPR/Cas9 genome editing approach. First, we designed a single guide RNA (sgRNA) targeting the *tgatp4* genomic locus near the region encoding amino acid residue 419. We introduced this sgRNA-encoding sequence into the pSAG1::Cas9-U6::sgUPRT plasmid (Addgene plasmid 54467) using Q5 mutagenesis (New England Biolabs), as described previously (56). For the Q5 mutagenesis, we used primers 12 and 13 (**Table S1**). We also generated a donor DNA consisting of the annealed primers 14 and 15 (**Table S1**), which encodes for a region homologous to the *tgatp4* locus that incorporates amino acid reside 419. To anneal these primers, we combined them at a 50 µM final concentration, heated them to 94°C in a heat block for 10 min, then allowed the heat block to cool to room temperature across several hours. We mixed the sgRNA-expressing plasmid, which also encodes Cas9-GFP, together with the donor DNA, and transfected the mixture into TgATP4-HA strain parasites, which were generated previously by fusing the TgATP4 protein to an HA-epitope tag in TATi/*Δku80* parasites (18). We selected and cloned GFP-expressing parasites two days after transfection using fluorescence activated cell sorting, as described previously (57). To identify parasite clones encoding the G419S mutation in the TgATP4 protein, we PCR amplified the *tgatp4* genomic locus from clonal parasite lines using primers 16 and 17 (**Table S1**), and subjected PCR products to Sanger sequencing. We identified multiple clones containing the G419S mutation (**Fig. S3**) and, following verification that the expression levels and localisations of the TgATP4^G419S^-HA protein did not differ between the various clones (**Fig. S4**), used clone B9 for subsequent experiments.

### Determination of the expression and localisation of TgATP4^G419S^

To compare the expression levels of the TgATP4^G419S^-HA protein to that of TgATP4^WT^-HA, we extracted proteins from the various parasite lines in LDS sample buffer (Thermo Fisher Scientific), loaded these at 2.5 × 10^6^ parasite equivalents per lane, and separated them by SDS-PAGE using Bis/Tris NuPAGE gels (12% acrylamide, Thermo Fisher Scientific) according to the manufacturer’s instructions. Samples were then transferred to nitrocellulose membranes using the Mini Blot module (Thermo Fisher Scientific), and subjected to western blotting. We probed blots with rat anti-HA (clone 3F10, Sigma catalogue number 11867423001) and anti-*Tg*Tom40 (58) primary antibodies, and horseradish peroxidase-conjugated goat anti-rat (Abcam, ab97057) and goat anti-rabbit (Abcam, ab97051) secondary antibodies. Blots were developed using a homemade chemiluminescence solution (0.04% w/v luminol, 0.007% w/v coumaric acid, 0.01% v/v H_2_O_2_, 100 mM Tris pH 9.4) and imaged using X-ray film.

To compare the localisation of the TgATP4^G419S^-HA protein to that of the TgATP4^WT^-HA protein, we undertook immunofluorescence assays as described previously (59). We probed samples with rat anti-HA (clone 3F10, Sigma catalogue number 11867423001) and mouse anti-*Tg*P30 (clone TP3, Abcam, ab8313) primary antibodies, and donkey anti-rat AlexaFluor 488 (Thermo Fisher Scientific, A-21208) and goat anti-rabbit AlexaFluor 546 (Thermo Fisher Scientific, A-11035) secondary antibodies. Samples were imaged using an Olympus IX71 microscope featuring a DeltaVision Elite set-up, with a 100X UPlanSApo objective lens (NA 1.40) and a CoolSNAP HQ2 camera. Images were deconvolved using SoftWoRx Suite 2.0 software, and adjusted linearly for contrast and brightness.

### Measurements of [Na^+^]_cyt_

Saponin-isolated mature trophozoite-stage *P. falciparum* parasites, or extracellular *T. gondii* tachyzoites, were loaded with the Na^+^-sensitive fluorescent dye SBFI (6 μM; in the presence of 0.01% w/v Pluronic F-127) for 20 min (*P. falciparum*) or ∼ 30 min (*T. gondii*), essentially as described previously (11,18). Measurements were performed essentially as described previously (11,18,19) with parasites suspended in pH 7.1 ‘Physiological Saline Solution’ (125 mM NaCl, 5 mM KCl, 1 mM MgCl_2_, 20 mM glucose and 25 mM HEPES). The ratio of the fluorescence intensity recorded at 340 nm and 380 nm (with an emission wavelength of 515 nm) was converted to [Na^+^]_cyt_ using a calibration procedure described previously (11). Fluorescence measurements and calibrations were performed at 37°C, either with individual 1 mL suspensions using a PerkinElmer LS 50B fluorescence spectrometer (for resting [Na^+^]_cyt_ measurements for which the data are shown in **Fig. S5**) or in 96 well plates using a Tecan Infinite M1000 PRO plate reader (all other [Na^+^]_cyt_ measurements).

### Measurements of membrane ATPase activity

Membranes from isolated *P. falciparum* or extracellular *T. gondii* parasites were prepared as described previously for *P. falciparum* (15), with protein concentrations in the membrane preparations measured using a Bradford assay (60). The PiColorLock Gold Phosphate Detection System (Innova Biosciences) was used to measure the production of P_i_ from hydrolysis of ATP, essentially as described previously (15). Membrane preparations were diluted in either a high Na^+^ solution (yielding final conditions in the reactions of 150 mM NaCl, 20 mM KCl, 2 mM MgCl_2_, 50 mM Tris, pH 7.2) or a Na^+^-free solution (yielding final conditions of 150 mM choline chloride, 20 mM KCl, 2 mM MgCl_2_, 50 mM Tris, pH 7.2) to achieve a total protein concentration of 50 μg/mL. Cipargamin, (+)-SJ733, or DMSO (solvent control) were added to give the concentrations stated in the relevant Figure legends. The reactions were performed at 37°C, and were initiated by the addition of 1 mM ATP (Na_2_ATP·3H_2_O; MP Biomedicals) and terminated 10 min later by adding 100 μL of reaction mixture (in duplicate) to 25 μL “Gold mix” in a 96 well plate. “Stabilizer” was added 3 min later, and the plates incubated at room temperature in the dark for 60 min before measuring absorbance at 635 nm. Background values (averaged from wells containing all components, but to which ATP was not added until after the membrane was exposed to Gold mix) were subtracted from the data. For each treatment, the ATPase activity associated with the (Na^+^-dependent) PfATP4 and TgATP4 proteins was calculated by subtracting data obtained in the low Na^+^ condition (containing only the 2 mM Na^+^ introduced on addition of 1 mM Na_2_ATP) from that obtained in the high Na^+^ condition.

### Comparisons of the growth rate of asexual *P. falciparum* parasites

The parasitaemias of synchronous cultures containing trophozoite-stage parasites were determined by flow cytometry. Briefly, *P. falciparum*-infected erythrocytes (1 mL) were centrifuged (2,000 × *g*, 30 s), and the cells were resuspended in 1 mL of pH 7.4 Physiological Saline Solution containing 20 µg/mL Hoechst 33258. The cells were incubated for 15 min at 37°C, and were then centrifuged and washed twice in pH 7.4 Physiological Saline Solution (2,000 × *g*, 30 s) before being resuspended in 1 mL of pH 7.4 Physiological Saline Solution. The cell suspension (50 μL) was then diluted into 250 μL pH 7.4 Physiological Saline Solution to a density of ∼ 10^6^ – 10^7^ cells/mL in 1.2 mL Costar polypropylene cluster tubes (Corning). Cells (100,000) were sampled from each tube at a low sampling speed with the following settings: forward scatter = 308 V (log scale), side scatter = 308 V (log scale), Alexa Fluor 488 = 559 V (log scale) and Pacific Blue = 478 V (log scale). A sample containing only uninfected erythrocytes was first analysed to establish a gating strategy that defined a threshold above which erythrocytes were deemed to be infected with *P. falciparum*. A sample containing wild-type *P. falciparum*-infected erythrocytes was then analysed to define a threshold below which parasites were deemed to be auto-fluorescent. Both gating strategies were then applied in all subsequent analyses to determine the parasitaemia.

The volume of saponin-isolated trophozoites was measured using a Beckman Coulter Multisizer 4 fitted with a 100 μm ‘aperture tube’, as described previously (12). The parasites were washed and resuspended (at 37°C) in pH 7.1 Physiological Saline Solution. The electrolyte solution within the aperture tube was also pH 7.1 Physiological Saline Solution. For each measurement of cell volume, approximately 20,000 pulses (each corresponding to the passage of a single cell through the aperture) were recorded. The mean volume of the parasites within each sample was determined by fitting a Gaussian distribution curve to the population data within a 6 – 60 fL window in GraphPad Prism 8.

## Acknowledgements

We are grateful to the Canberra Branch of the Australian Red Cross Lifeblood for the provision of blood, to the Medicines for Malaria Venture for the provision of cipargamin and (+)-SJ733, and to Assoc. Prof. Erkang Fan and Prof. Akhil Vaidya for the provision of PA21A050. We also thank Prof. Elizabeth Winzeler for the Dd2-PfATP4^T418N,P990R^ and Dd2 parental lines, and Prof. R. Kip Guy for (+)-SJ733 and for the W2-PfATP4^P966S^, W2-PfATP4^P966T^ and W2 parental lines. We are grateful to Xinxin Zhang, Sasha Lee and Kwong Sum Shea for initial exploratory Na^+^ assays and ATPase assays with the HCR and Dd2-PfATP4^T418N,P990R^ lines, and to Jennifer Thompson and Prof. Alan Cowman for assistance with whole genome sequencing.

