## Supplemental Table, Data and Figures for "Generation and characterisation of *P. falciparum* parasites with a G358S mutation in the PfATP4 Na^+^ pump and clinically relevant levels of resistance to some PfATP4 inhibitors"

**Table S1. Primers used for *pfatp4* amplification and sequencing in this study, and for the genetic modification of *T. gondii*.**

| Primer | Sequence (5' – 3') | Use | Reference |
|---|---|---|---|
| ***P. falciparum*** | | | |
| **1** | ATGAGTTCTCAAAATAATAATAAACAGG | Amplification and sequencing | (1) |
| **2** | TTAATTCTTAATAGTCATATATTTTCTTCTATATATAACC | Amplification and sequencing | (1) |
| **3** | TCACCACAATGTACTGTGTTAAGAAA | Sequencing | (1) |
| **4** | ATCCAACAAAAGGTTTTGAACCATGT | Sequencing | (1) |
| **5** | TACAAACATGTAGAGAAGCACAAGTT | Sequencing | (1) |
| **6** | TATCTCCGTCTTCTACATTATTG | Sequencing | (2) |
| **7** | ATGACAGCAATTAATGCAGTTAC | Sequencing | (2) |
| **8** | CATGTAGTATATCTGCAACTTTAAC | Sequencing | (2) |
| **9** | ATATGAGTGAAGGACCAAT | Sequencing | |
| **10** | CTTTAGCTATAGAATGTTCA | Sequencing | |
| ***T. gondii*** | | | |
| **11** | CTGTAGTAGTTATGAATATA | Sequencing | |
| **12** | CAACGTGGCCTGAATCGACTGTTTTAGAGCTAGAAATAGCAAG | Q5 mutagenesis | |
| **13** | AACTTGACATCCCCATTTAC | Q5 mutagenesis | |
| **14** | AGGCTCCAGGCTCACTCCCCTTCAACGTGGCCTGAATCGACTG TCTGGCATGATTGGACTGATCGCCATCTGCGTCCTCATCATCGT CGT | Annealing | |
| **15** | ACGACGATGATGAGGACGCAGATGGCGATCAGTCCAATCAT GCCAGACAGTCGATTCAGGCCACGTTGAAGGGGAGTGAGCC TGGAGCCT | Annealing | |
| **16** | GGTCGAACTGAAGACGAACG | Sequencing | |
| **17** | GTTTGAGCGTACAGTGAAGACG | Sequencing | |

**Data S1. Whole genome sequencing results for HCR1 and HCR2 parasites, their Dd2-PfATP4$^{T418N,P990R}$ parent, and its Dd2 parent.**

Four clonal samples sequenced for this study:

Dd2 (parent of Dd2-PfATP4$^{T418N,P990R}$)

Dd2-PfATP4$^{T418N,P990R}$ (parent of HCR1 and HCR2)

HCR1 (Highly Cipargamin Resistant clone 1)

HCR2 (Highly Cipargamin Resistant clone 2)

| Sample | coverage when aligned to Dd2 reference | coverage when aligned with 3D7 reference |
|---|---|---|
| Dd2 | 212.8 | 229.5 |
| Dd2-PfATP4$^{T418N,P990R}$ | 192.9 | 208.2 |
| HCR1 | 118.6 | 129.7 |
| HCR2 | 228.9 | 249.0 |

Dd2 alignment rates: 86% - 87%; 3D7 alignment rates: 98%

All subsequent bioinformatics analysis uses reads aligned to PlasmoDB-29_Pfalciparum3D7 reference genome.

Copy Number Analysis was performed using QDNAseq with 5 and 1 kbp bins using the recommended workflow. Binned counts were filtered by min-mappability=50%, and a blacklist of centromere and telomere regions, then a loess fit used to correct for GC and mappability, and counts converted to copy numbers. They were inspected as copy numbers and as scaled relative to the parent strain.

Dd2-PfATP4$^{T418N,P990R}$ and Dd2 both had the same amplification on chromosome 5. Scaled, relative copy numbers showed no features of interest. HCR1 and HCR2 copy numbers scaled relative to Dd2-PfATP4$^{T418N,P990R}$ both showed a duplication on chromosome 12, and HCR2 had a relative deletion on chromosome 5, showing a reduction in the observed amplification (**Fig. S1**).

Structural variation analysis used GRIDSS, with candidate SVs inspected in Integrated Genome Viewer (IGV), found no new structural variations in Dd2-PfATP4$^{T418N,P990R}$ compared to Dd2. There was a clear event in chromosome 12 in both HCR1 and HCR2 (**Fig. S1**). This is a duplication from 520kb – 556kb, covering nine genes including

PF3D7_1211900, which is *pfatp4*. The breakends are in homopolymer runs of A, at Pf3D7_12_v3:520014..520039 and Pf3D7_12_v3:556464..556489.

Single-nucleotide changes and small insertions and deletions were called with SNVer. VarScan was also run but did not give usable results. All calls which also appeared in the parent strain were discarded, and remaining calls filtered by depth at least 10, alternate-frequency > 0.4, discard calls in the first and last 10% of each chromosome. When calls are filtered for 'in a coding sequence', ignore genes described as PfEMP1, rifin, stevor or pseudogene. This left 100s of SNPs and 10s of indels. On inspection most of these events were clustered in areas of low coverage, or highly repetitive regions, and some of the remainder were synonymous changes.

### Small variants of possible interest

| Sample | Position | AltCount/ Depth | Change | Gene | Comment or description |
|---|---|---|---|---|---|
| Dd2-PfATP4$^{T418N, P990R}$ | chr12: 529831 | 108/108 | G->C; P->R | PF3D7_1211900 | Known P990R in PfATP4 |
| Dd2-PfATP4$^{T418N, P990R}$ | chr12: 531547 | 102/102 | G->T; T->N | PF3D7_1211900 | Known T418N in PfATP4 |
| HCR1, HCR2 | chr12: 531728 | 73/146 and 158/342 | C -> T; G->S | PF3D7_1211900 | G358S, with AF=0.5 |
| HCR1, HCR2 | chr5: 851823 | 103/103 and 246/246 | T->C; M->V | PF3D7_0520800 | |
| HCR1, HCR2 | chr7: 605674 | 9/10 and 22/22 | G->A; V->I | PF3D7_0713100 | Depth is borderline. Pfmc-2TM: Maurer's |
| HCR1, HCR2 | chr10: 1439013 | 41/41 and 88/88 | G->A; V->I | PF3D7_1036400 | liver: stage |
| HCR2 | chr3: 136831 | 53/57 | C -> G; Q->E | PF3D7_0302500 | cytoadherence |

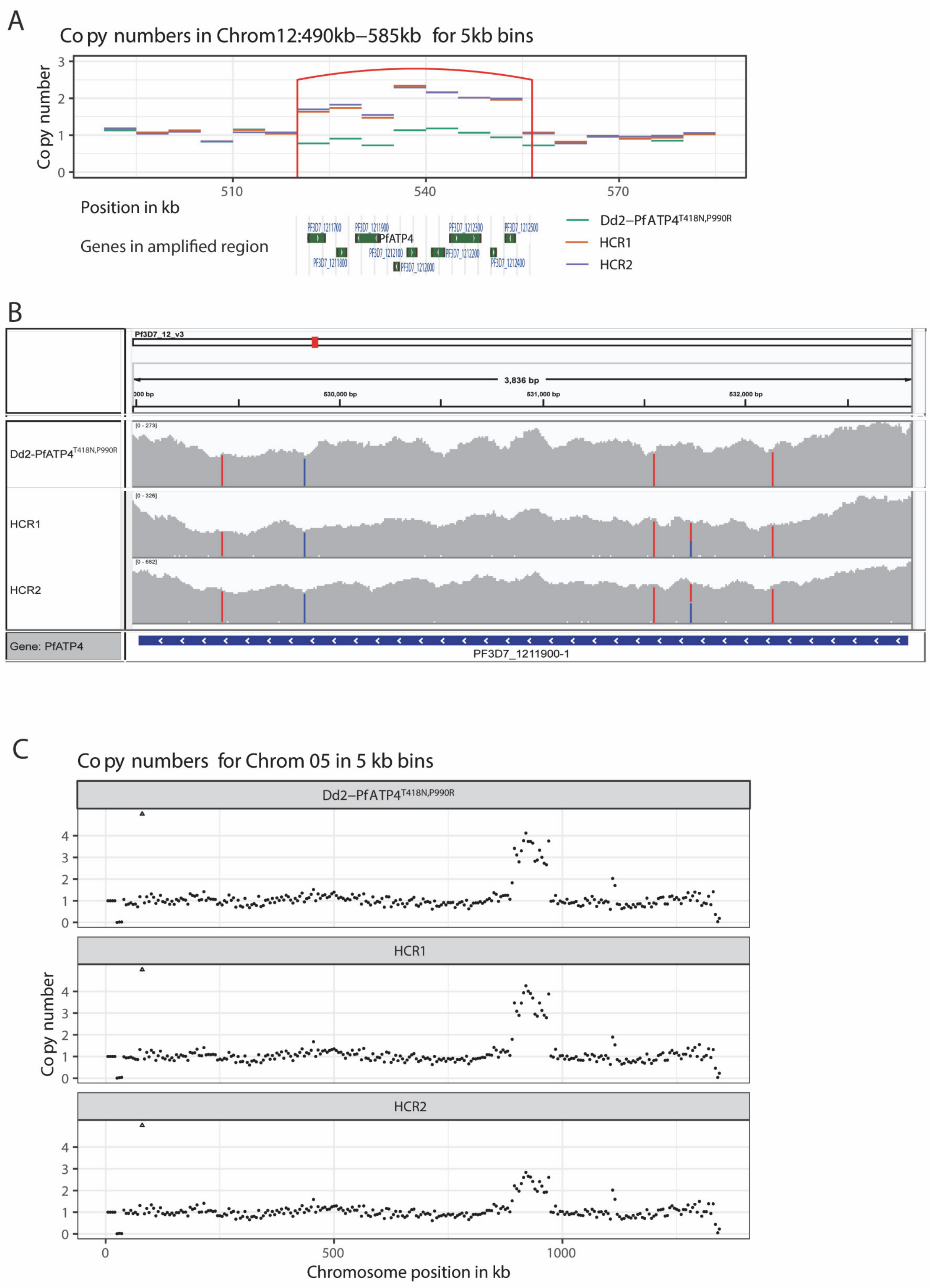


**Fig. S1. Whole genome sequencing results for Dd2-PfATP4$^{T418N,P990R}$, HCR1 and HCR2. A**. Duplicated region in chromosome 12 present in HCR1 and HCR2. Red vertical lines indicate breakpoint boundaries. **B**. Integrated Genome Viewer image of PfATP4 region. G358S is in HCR1 and HCR2 with allele frequency of 0.5. The mutations giving rise to T418N and P990R are shown. Two mutations found in Dd2 parasites relative to 3D7 parasites (one synonymous and one giving rise to a G1128R change) are also shown. Bars are coloured by nucleotide if more than 25% of reads differ from reference. Red = T, blue = C. **C**. Differently-amplified region in chromosome 5, also detected by structural variant caller. The break ends are at 888 kb and 970 kb, including gene *pfmdr1*, location 955,955..963,095(+). Three or four copies are present in Dd2- PfATP4$^{T418N,P990R}$ and HCR1, and one fewer in HCR2.

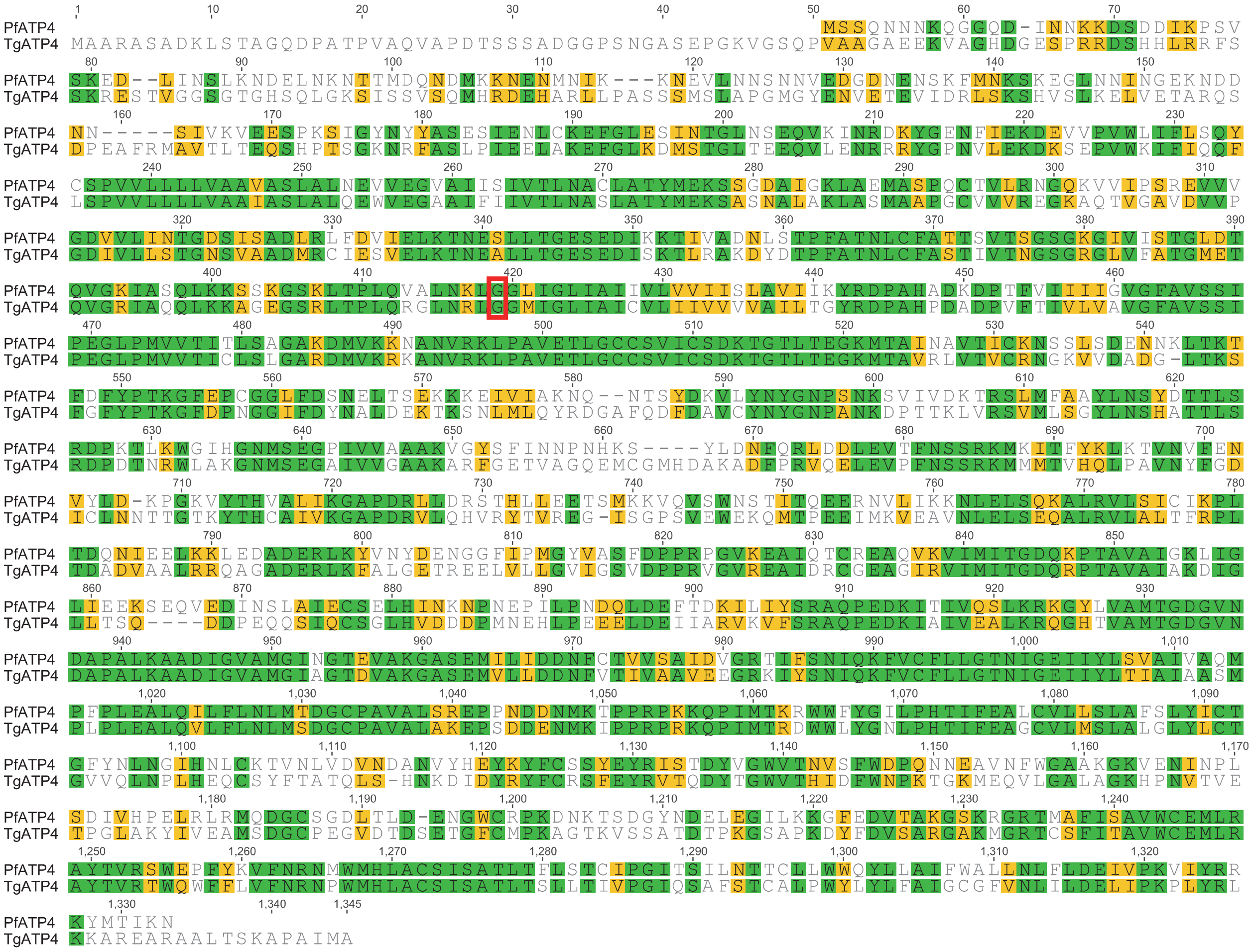


**Fig. S2. Sequence alignment of PfATP4 and TgATP4 and location of the G358 (PfATP4) and G419 (TgATP4) residues.** The protein sequences for PfATP4 (PF3D7_1211900) and TgATP4 (TGGT1_278660) were aligned in Geneious. Identical residues are shown in green and residues with similar chemical characteristics are shown in yellow. The G358 and G419 residues are shown in a red box.

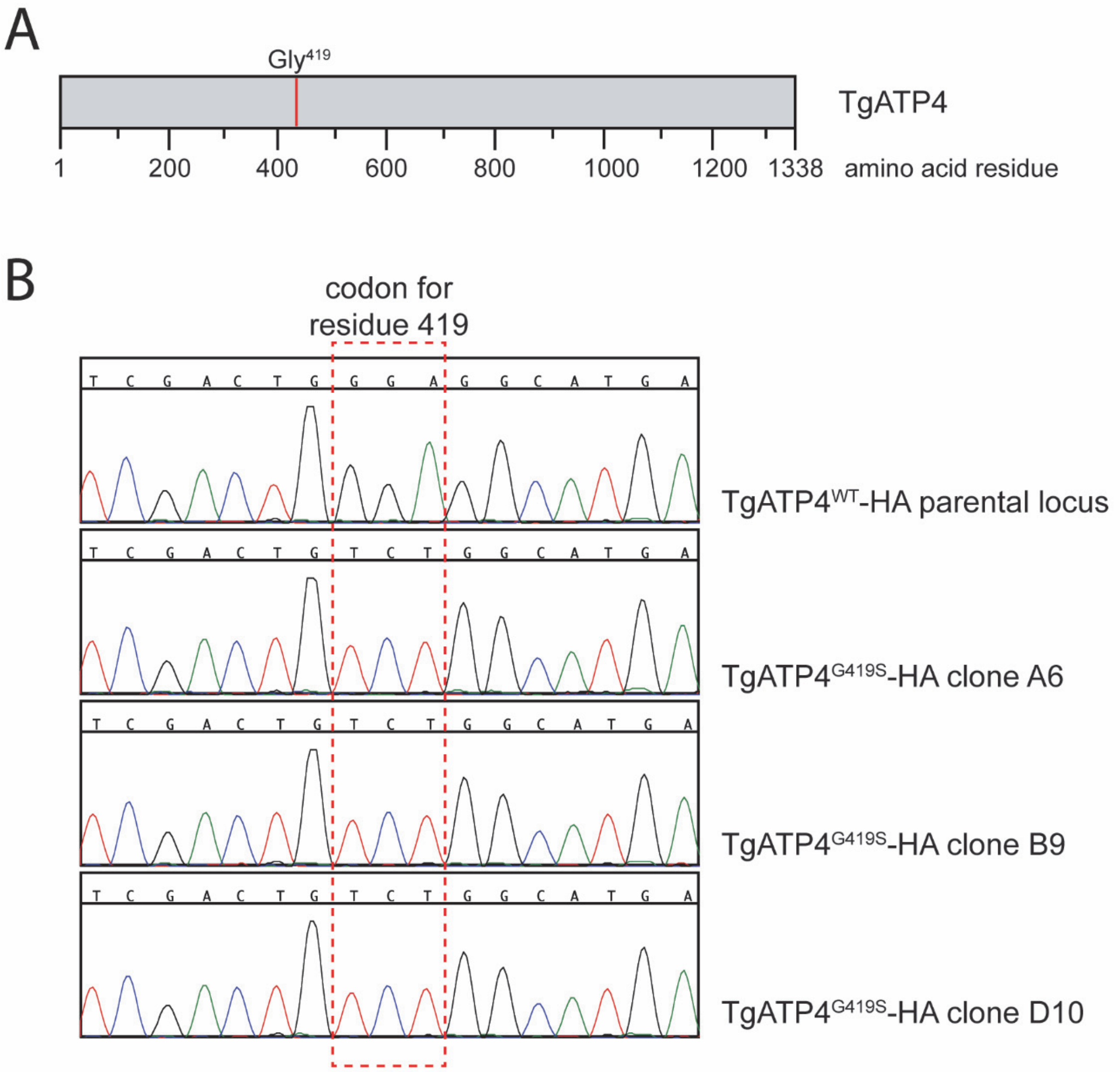


**Figure S3. Generation of TgATP4$^{G419S}$-HA-expressing *T. gondii* parasite clones. A.** Schematic of the amino acid sequence of TgATP4, drawn to scale and depicting the position of the glycine residue at amino acid position 419 (Gly$^{419}$). **B.** Sanger sequencing chromatograms of the region of the *tgatp4* genomic locus that encodes residue 419 in a parasite clone containing the 'wild type' parental locus (TgATP4$^{WT}$-HA; top) and in parasite clones containing the G419S-encoding mutations (TgATP4$^{G419S}$-HA, clones A6, B9 and D10). The red box depicts the codon that encodes residue 419, with a glycine (GGA) encoded in the parental strain, and a serine (TCT) encoded in the modified TgATP4$^{G419S}$-HA clones.

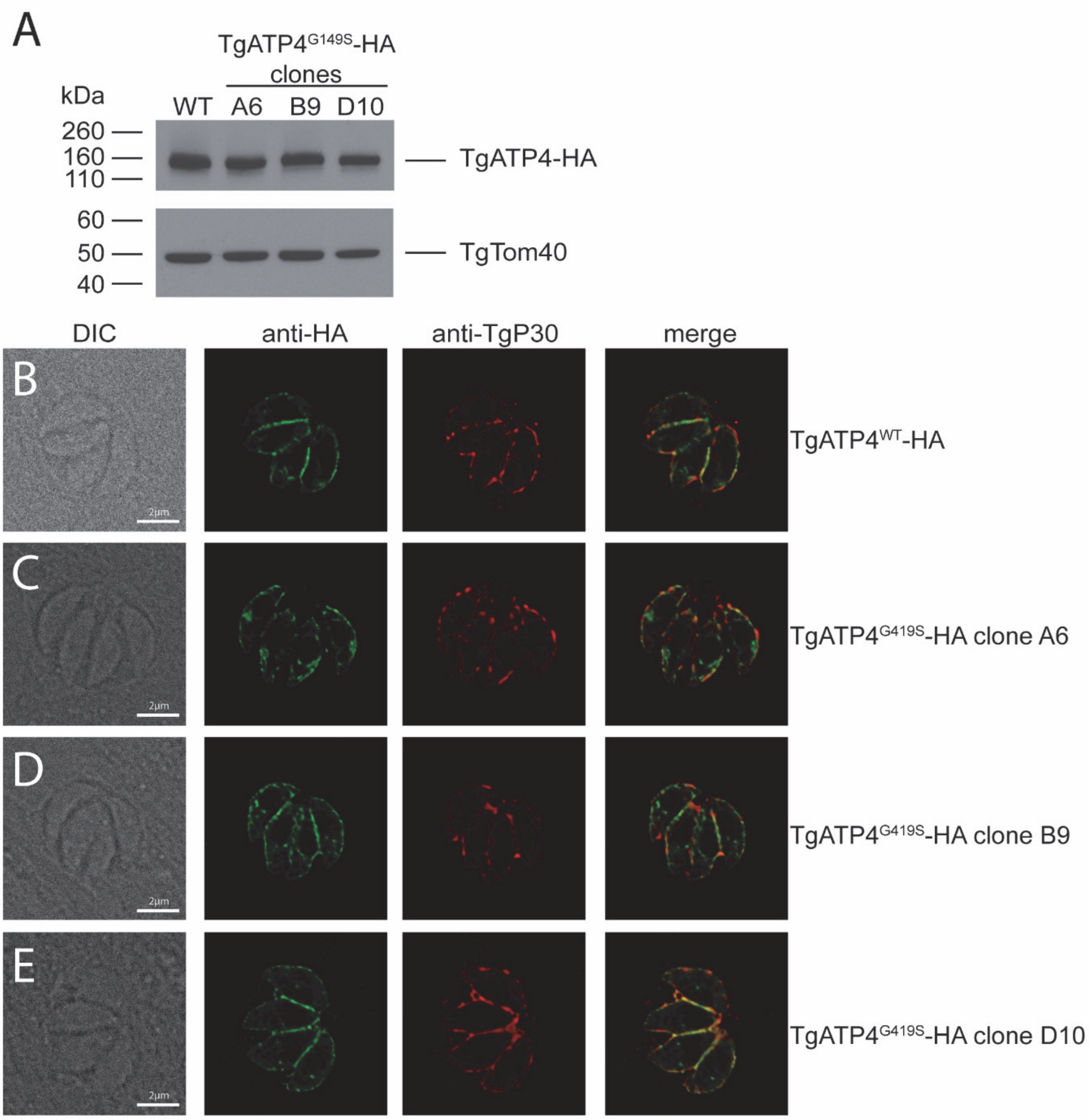


**Figure S4. Expression and localisation of TgATP4$^{G419S}$-HA in *T. gondii* parasites. A.** Western blot of TgATP4$^{WT}$-HA and TgATP4$^{G419S}$-HA expressing parasites, probed with anti-HA antibodies (top) and anti-TgTom40 antibodies as a loading control (bottom). **B-E.** Immunofluorescence assays of TgATP4$^{WT}$-HA (**B**) and TgATP4$^{G419S}$-HA expressing parasite clones (**C-E**) probed with anti-HA antibodies (green) and anti-*Tg*P30 antibodies as a marker for the plasma membrane (red). Scale bars are 2 µm. DIC, differential interference contrast transmission images.

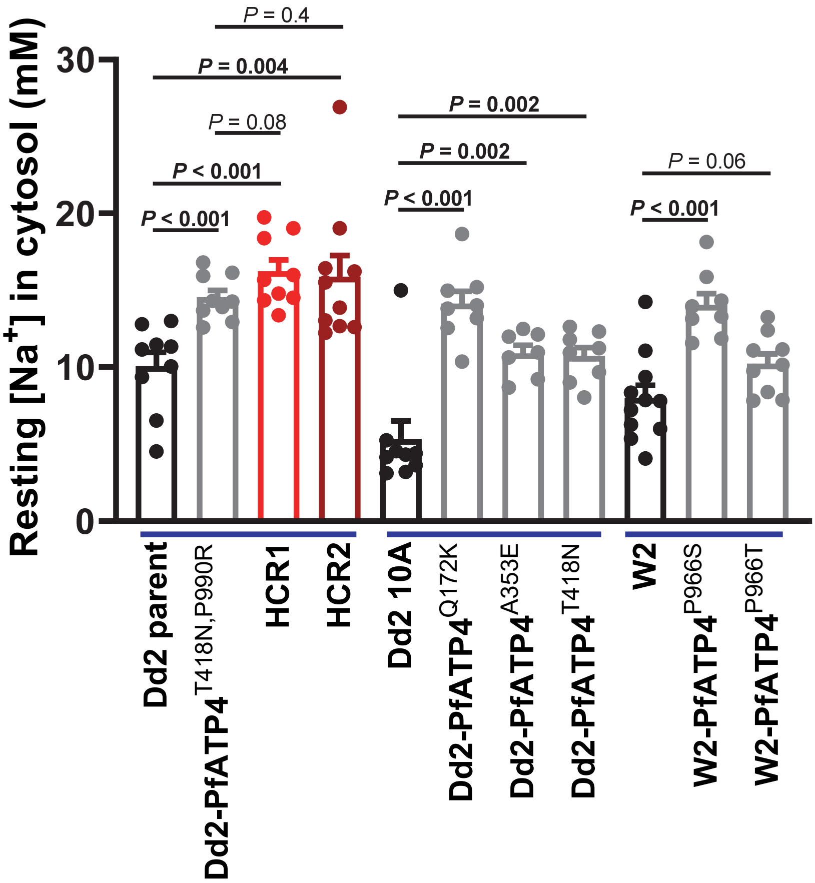


**Fig. S5. Resting cytosolic [$Na^+$] for parasite lines expressing wild-type or mutant variants of PfATP4.** Data for three sets of lines are shown (see Methods for details of their origins), with the data for parental lines shown in black, those for parasites with low-level cipargamin resistance shown in grey, and those for parasites with high-level cipargamin resistance shown in red. The measurements were performed with isolated trophozoite-stage parasites loaded with the $Na^+$-sensitive dye SBFI, and suspended in Physiological Saline Solution (pH 7.1) at 37˚C. The bars show the mean + SEM obtained from at least seven independent experiments for each line, and the symbols show the results obtained in individual experiments. The *P* values shown are from unpaired t-tests (with significant differences shown in bold).

1. Rottmann, M., McNamara, C., Yeung, B. K., Lee, M. C., Zou, B., Russell, B., Seitz, P., Plouffe, D. M., Dharia, N. V., Tan, J., Cohen, S. B., Spencer, K. R., Gonzalez-Paez, G. E., Lakshminarayana, S. B., Goh, A., Suwanarusk, R., Jegla, T., Schmitt, E. K., Beck, H. P., Brun, R., Nosten, F., Renia, L., Dartois, V., Keller, T. H., Fidock, D. A., Winzeler, E. A., and Diagana, T. T. (2010) Spiroindolones, a potent compound class for the treatment of malaria. *Science* **329**, 1175-1180
2. Lehane, A. M., Ridgway, M. C., Baker, E., and Kirk, K. (2014) Diverse chemotypes disrupt ion homeostasis in the malaria parasite. *Mol Microbiol* **94**, 327-339
